# The spiking output of the mouse olfactory bulb encodes large-scale temporal features of natural odor environments

**DOI:** 10.1101/2024.03.01.582978

**Authors:** Suzanne M. Lewis, Lucas M. Suarez, Nicola Rigolli, Kevin M. Franks, Nicholas A. Steinmetz, David H. Gire

## Abstract

In natural odor environments, odor travels in plumes. Odor concentration dynamics change in characteristic ways across the width and length of a plume. Thus, spatiotemporal dynamics of plumes have informative features for animals navigating to an odor source. Population activity in the olfactory bulb (OB) has been shown to follow odor concentration across plumes to a moderate degree (Lewis et al., 2021). However, it is unknown whether the ability to follow plume dynamics is driven by individual cells or whether it emerges at the population level. Previous research has explored the responses of individual OB cells to isolated features of plumes, but it is difficult to adequately sample the full feature space of plumes as it is still undetermined which features navigating mice employ during olfactory guided search. Here we released odor from an upwind odor source and simultaneously recorded both odor concentration dynamics and cellular response dynamics in awake, head-fixed mice. We found that longer timescale features of odor concentration dynamics were encoded at both the cellular and population level. At the cellular level, responses were elicited at the beginning of the plume for each trial, signaling plume onset. Plumes with high odor concentration elicited responses at the end of the plume, signaling plume offset. Although cellular level tracking of plume dynamics was observed to be weak, we found that at the population level, OB activity distinguished whiffs and blanks (accurately detected odor presence versus absence) throughout the duration of a plume. Even ∼20 OB cells were enough to accurately discern odor presence throughout a plume. Our findings indicate that the full range of odor concentration dynamics and high frequency fluctuations are not encoded by OB spiking activity. Instead, relatively lower-frequency temporal features of plumes, such as plume onset, plume offset, whiffs, and blanks, are represented in the OB.

## Introduction

Brains generate complex and adaptive behavioral sequences (Dennis et al., 2021; Miller et al., 2022), a subset of which are driven by olfactory stimuli and are important for navigation and the search for resources (Baker et al., 2018). Theoretical work has suggested that the spatiotemporal dynamics of natural odor environments could enable multiple types of search strategies that would allow for successful olfactory-guided navigation. (Boie et al., 2018; Gardiner & Atema, 2010; Gumaste et al., 2020; P. W. Jones & Urban, 2018; Loisy & Eloy, 2022; Park et al., 2016; Rigolli et al., 2022; Vergassola et al., 2007; Vickers, 2000). Experimentally, recent work has focused on generating a better understanding of which strategies navigating mice use (Findley et al., 2021; Gire et al., 2016) to enable their repertoire of complex odor-driven behaviors. These strategies are constrained by the features of odor cues that are encoded by the brain. Olfactory cortical neurons respond to isolated features of natural odor dynamics (Ackels et al., 2021; Dasgupta et al., 2022; Lewis et al., 2021; Parabucki et al., 2019), but the neural substrates that enable odor-driven search in mice are not yet known.

To understand the ease with which mice navigate plumes despite the complexity of plume dynamics, we must understand how plume dynamics inform olfactory search and what subset of dynamical features are encoded by olfactory processing.

A navigating mouse encounters odors as plumes, which are the result of the turbulent mixing of air and odor particles (Celani et al., 2014). As odor moves away from a source, it is pulled and pushed into discrete filaments. Thus, plumes are necessarily encountered as “whiffs” of odor separated by “blanks” of pure air (Riffell et al., 2008), and the properties of odor encounters show characteristic changes across the length and width of a plume (Celani et al., 2014; Mylne & Mason, 1991). Since whiff properties, such as how often they are encountered (Murlis et al., 1992; Mylne & Mason, 1991) and how long they last (Young et al., 2020; Rigolli et al., 2022), have structure across plume space, they contain information regarding an animal’s position relative to an odor source.

Olfactory processing occurs in a relatively shallow system as compared to other senses with no relay through the thalamus (Shepherd, 2005). This means that all ethologically-relevant odor information must be conveyed by the olfactory bulb (OB), the first olfactory relay of the brain and the last common structure of the olfactory pathway (Mori et al., 1999). Single cells in the OB accurately resolve odor presented in isolated pulses or isolated frequencies (Ackels et al., 2021; Dasgupta et al., 2022), but little is known about whether the OB follows stochastic plume dynamics, with mixed frequencies. Additionally, there are known adaptation effects in the OB (Martelli & Storace, 2021) and dynamical changes in OB responses both over the course of longer odor stimuli (Baker et al., 2019; Patterson et al., 2013; Spors & Grinvald, 2002) or over back to back sampling across sniffs (Fukunaga et al., 2012). Little is known regarding whether the dynamics of adaptation or the dynamics of OB responses themselves may compromise the ability of the OB to detect odor with naturalistic and complex dynamics across longer timescales. The population activity of mitral and tufted cells (MTCs) in the OB respond to stochastic plumes at the glomerular level as measured by widefield calcium imaging (Lewis et al., 2021). The collective MTC activity across dorsal OB imaging windows was analyzed using principal component analysis, and each field of view had a high-ranking principal component that was moderately correlated with concentration dynamics across the plume (∼0.6 average Pearson correlation). Segmenting the population into individual putative glomeruli, spherical complexes where olfactory receptor neurons that express the same receptor type synapse onto OB cells, showed relatively weaker but still moderate tracking ability among glomeruli. Glomeruli varied in their ability to follow plume dynamics, but the best tracking glomeruli exhibited moderate correlations with plume dynamics across plumes (∼0.4 Pearson correlation). Therefore, at the population level, plume dynamics structure activity in the OB, but how individual cells or cell types contribute to the following behavior observed in collective MTC activity could not be determined by the widefield imaging approach.

Two main classes of OB projection neurons, mitral cells (MC) and tufted cells (TC), receive odor input from olfactory sensory neurons within glomeruli. MCs and TCs differ in their laminar location within the OB (Fukunaga et al., 2012; Igarashi et al., 2012), and the nature of the inhibitory surround that impinges upon them (S. Jones et al., 2020). Additionally, the two cells types differ in the nature of their input. Tufted cells receive more direct sensory input than MCs (Fukunaga et al., 2012; Gire et al., 2012; Gire & Schoppa, 2009; Najac, Jan, et al., 2011). MCs receive more indirect sensort input through their extensive connections with other MCs. This is a result of the dendrodendritic connections MCs form with granules cells. These connections also result in a greater amount of surround inhibition for MCs (Geramita et al., 2016), exerted upon their millimeter-long lateral dendrites (Aghvami et al., 2022). This could possibly provide individual MCs with informative lateral inhibition from other MCs across the bulb. These differences between MCs and TCs have led to the hypothesis that the two cell types play different roles in encoding odor information (Nagayama et al., 2004).

In this current study, we used high-density multi-shank Neuropixels 2.0 (Steinmetz et al., 2021) probes to simultaneously record from large populations of MTCs responding to plume dynamics at both the individual and population level in awake, head-fixed mice. We released odor from an upwind odor source and simultaneously recorded both odor concentration dynamics and cellular responses. We used previously-reported local field potential (LFP) signatures of laminar layers to separate cells into putative MCs and more superficial cells (EPL/SFL cells, predominantly putative TC). To capture odor concentration dynamics across plumes we used an adapted, miniature odor sensor (Tariq et al., 2021) to directly recorded the plume signal at the mouse nose. We then compared this sensory input with both individual cell spiking and population activity. We found that OB cell spiking encodes large-scale temporal features of odor plumes. Here we use large-scale in a relative manner, as plumes contain both high frequency and low frequency spatiotemporal features. OB activity resolved plume onset, plume offset, and whiffs and blanks, which are all lower-frequency (∼4Hz or lower) and binarized features of plumes (Celani et al., 2014). At the individual level, OB cells signaled moments when animals encountered or lost a plume. At the population level, the OB discerned odor presence accurately across the duration of the plume. Despite previous widefield findings that large OB populations can collectively track odor concentration with moderate strength, at a more local scale, cells were not observed to tightly track the full dynamics of odor concentration fluctuations. Instead, the majority of OB activity follows plume dynamics in the form of large-scale temporal features such as plume onset, plume offset, and whiff encounters, features thought to be vital for odor-guided navigation (Park et al., 2016; Vickers, 2000) and source localization (Rigolli et al., 2022).

## Results

### Physiological features of cells with relation to their distance from the MCL

High-density recordings from the OB of awake-headed mice during plume presentations was acquired using multi-shank Neuropixels 2.0 probes (Fig 1). Mice were head-fixed within a wind tunnel while odor was released from an upwind odor source (see methods). The same odor mixture was released each trial, but as plume dynamics are stochastic, odor concentration time series differed for every trial (Fig 1d). We recorded the odor concentration changes across the plume using a modified, odor sensor (Tariq et al., 2021) while simultaneously measuring plume-elicited responses in the OB. Before analyzing neural responses to plume dynamics, we first sought to divide cell types by estimating the position of MCL.

**Figure 1:**
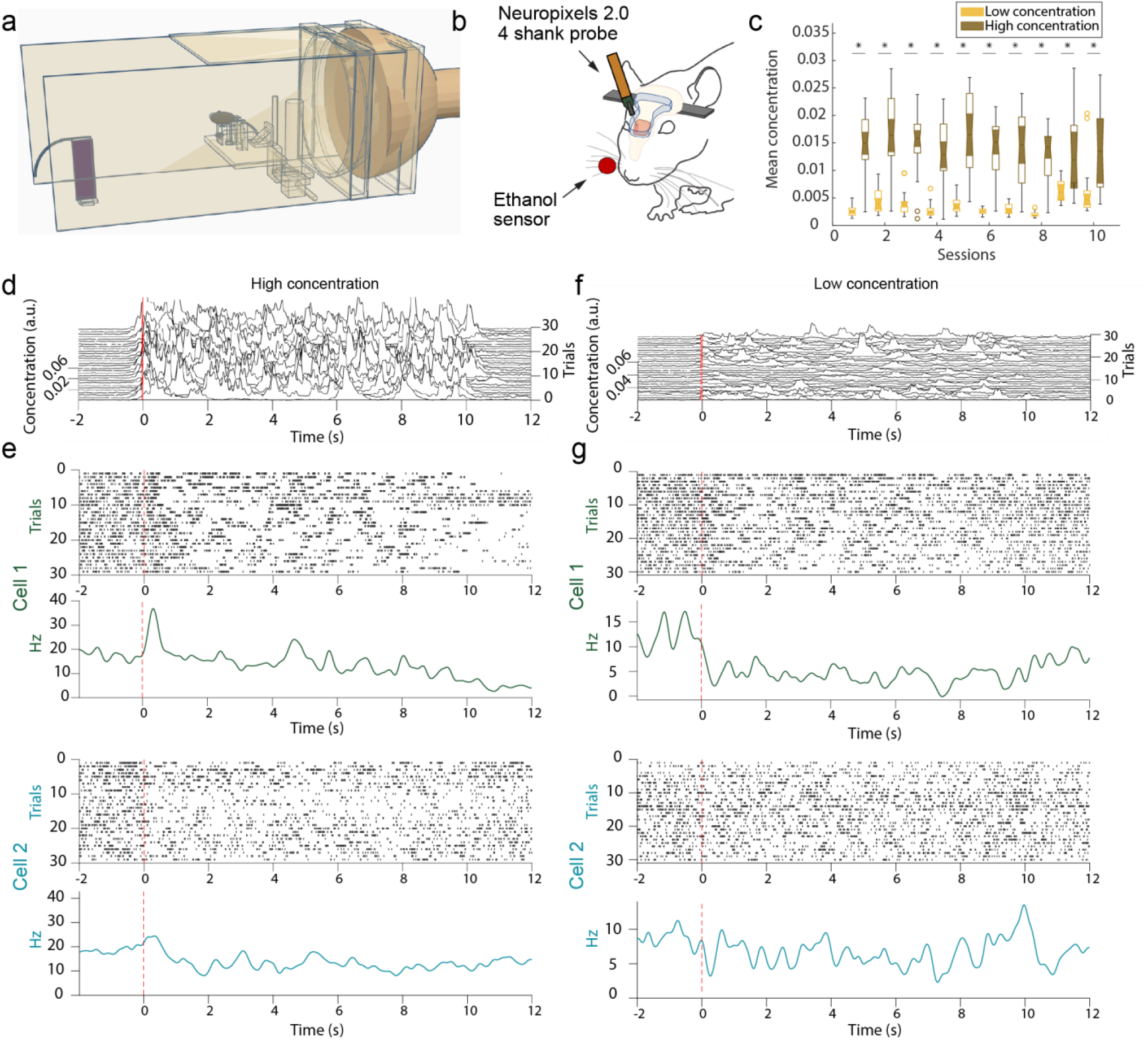
Experimental setup for acquisition of high-desnity electrophysiological recordings using Neuropixels 2.0 electrode arrays in the OB. **(A)** The wind tunnel in which plumes released by an upwind odor source (left) are pulled past the mouse (middle) by a rear exhaust vacuum. **(B)**. Neural population activity in the dorsal olfactory bulb was recorded using 4 shank, Neuropixels 2.0 probes. Odor concentration was recorded using a modified, passive ethanol sensor placed ∼4mm from the edge of the nostril closest to the sensor. **(C)** Plumes of high concentration (brown) and low concentration (yellow) were presented (see methods). The average concentration between low flow (high concentration) and high flow (low concentration) trials is significantly different for each session. **(D)** Plume dynamics were stochastic resulting in unique odor concentration dynamics for each trial. Odor concentration is plotted for all high concentration (low flow) trials in a single session aligned to the onset of the first whiff of each plume (red). **(E)** Example responses from two cells across high concentration trials are plotted beneath. Raster plots (top) and PSTHs (bottom) aligned to plume onset (dashed red line) exhibit different responses between the two cells to both the plume and the offset of the plume (∼10 s). **(F-G)** Same as (d-e) but for low concentration (high flow) trials.

Known characteristics of LFP across the laminar layers of the bulb was used to estimate the location of the MCL (see methods). We used a continuous wavelet transform approach to estimate LFP at gamma (30-100 Hz), theta (2-10 Hz), and spiking (300+ Hz, where 434Hz is the upper limit of the down sampled LFP) frequencies (see methods). We then manually located both polarity reversal in gamma and theta power and a sharp drop off in spiking power to estimate the position of MCL along the probe (Fig 2). In a single shank Neuropixels 2.0 recording that traversed the dorsal to ventral length of the OB, we found two striking examples, and only two examples, of these coincident changes in gamma, theta, and spiking power (Extended Data Fig 1), allowing localization of the dorsal and ventral intersections of the MCL with the probe. Using this approach, LFP characteristics were used to estimate the location of MCL on each shank of the four shank Neuropixels 2.0 probe.

**Figure 2:**
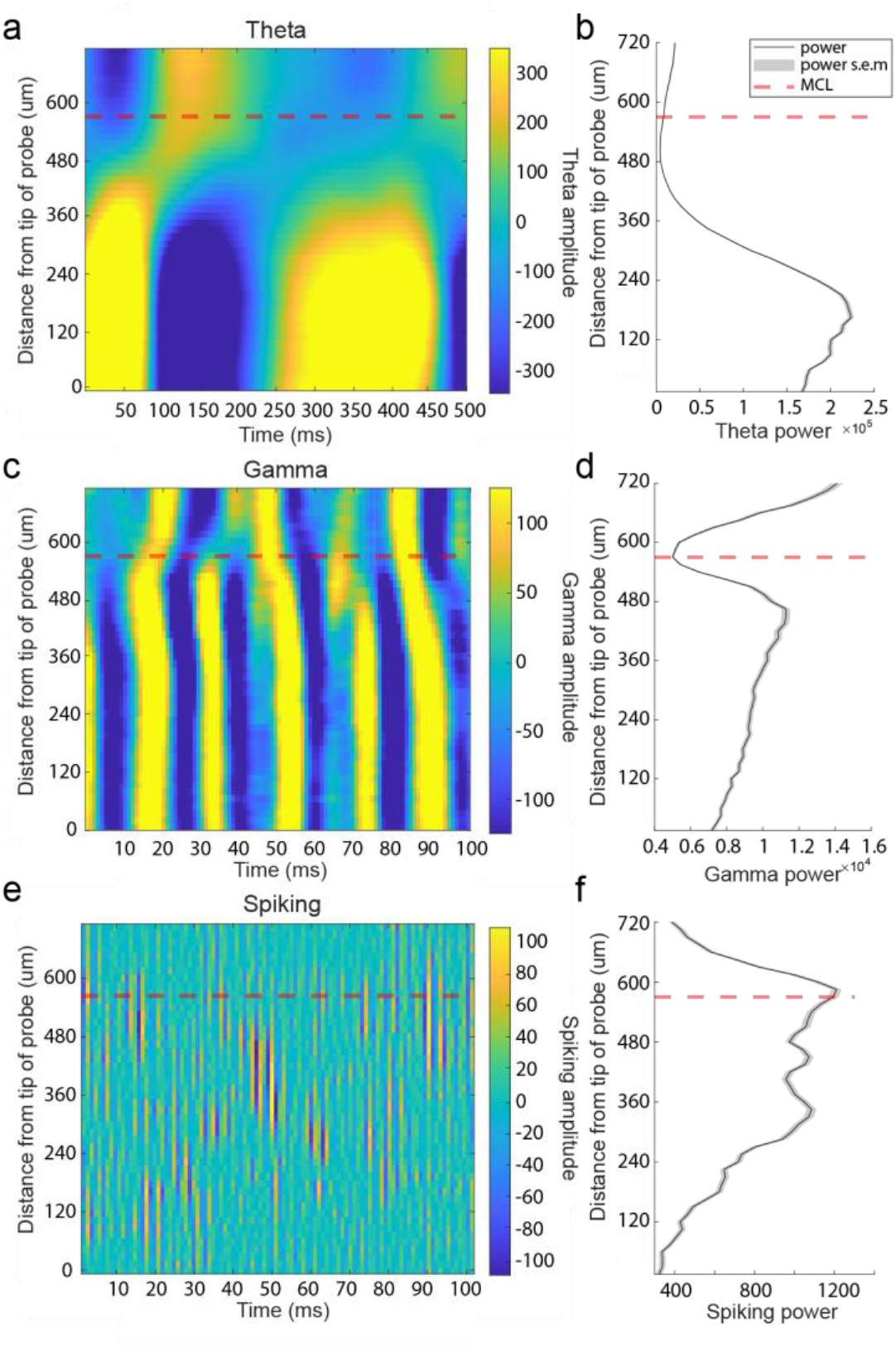
Local field potential at theta, gamma, and spiking frequencies was used to estimate the position of the MCL along recording arrays. **(A)** Theta LFP (2-10 Hz) amplitude plotted across 500 ms from recording sites across 1 of the probe’s 4 shanks from a single recording session of the ventral OB. Theta LFP, extracted using CWT based method, exhibits polarity reversal in the vicinity of the estimated MCL (red dashed line). Recording sites below the MCL are more superficial sites located closer to the ventral surface of the OB. **(B)** Plot of average theta power across 5 seconds of spontaneous activity for each recording site. **(C-D)** Same as (a-b) but for gamma frequency band amplitude (30-100 Hz) showing a drop in gamma amplitude near the MCL and, in this case, a reversal of amplitude polarity on either side of the MCL. **(E-F)** Same but for spiking frequencies (300+ Hz) showing most of the spiking amplitude decreases rapidly when moving away from the MCL towards the IPL/GCL (recording sites plotted above the MCL).

After estimating the location of the MCL (Extended Data Fig 2), the distance of each row of electrode sites from the center of the MCL was calculated. Cell layers were then defined based on their position relative to the estimated MCL. Across all recordings and shanks for which there were LFP characteristics of an MCL, we recorded 100 cells in the MCL, 218 cells in the EPL/SFL (external plexiform layer/ more superficial layers), and 147 clusters in the IPL/GCL (internal plexiform layer/ granule cell layer) (Extended Data Fig 3). Representative waveforms (Fig 3b-i) show cells with high amplitude waveforms in the principal cell layers. Waveform amplitude distributions were overlapping between areas (Fig 3j), but mean amplitude was highest in the MCL (m = 297.00 µv, std = 145.15 µv). It was significantly higher than the mean waveform amplitude of cells in the EPL/SFL (m = 231.48 µv, std = 114.42 µv) (Z = 4.03, p < 0.001, d = 0.53) and in the IPL/GCL (m = 126.24 µv, std = 77.56 µv) (z = 10.05, p<0.001, d=1.55). Importantly, mean waveform amplitude in the IPL/GPL was significantly lower as compared to either other area (Table 2). Therefore, waveform amplitude is highest in the MCL, second highest in the EPL/SFL, and lowest in the GCL. This alignment also reflects the fact that mean soma sizes for each area also decrease in size in a similar manner. MC somas are on average larger than most tufted cell (TC) subtypes in the EPL/SFL (Nagayama et al., 2010), and average GCs soma size is the smallest mean soma size across OB layers (Nagayama et al., 2014).

**Table 1:** Comparing physiological features relative to the estimated MCL. Significant changes in both waveform amplitude and mean spiking rate were observed between indicated areas. Difference in physiological features between areas were calculated using a Wilcoxon rank-sum test, and effect size was calculated using Cohen’s d. Cells or clusters located on an electrode array that did not have an identifiable MCL were excluded from the analysis.

|  | Mean | Standard deviation | n | Combination | rank-sum score | p-value | cohen's d |
| --- | --- | --- | --- | --- | --- | --- | --- |
| Waveform amplitude |  |  |  |  |  |  |  |
| EPL/SFL | 231.48 | 114.42 | 218 | EPL/SFL & IPL/GCL | 9.72 | p<0.001 | 1.04 |
| MCL | 297.00 | 145.15 | 100 | MCL & EPL/SFL | 4.03 | p<0.001 | 0.52 |
| IPL/GCL | 126.24 | 77.56 | 147 | MCL & IPL/GCL | 10.05 | p<0.001 | 1.55 |
| Spiking frequency |  |  |  |  |  |  |  |
| EPL/SFL | 14.03 | 9.05 | 18 | EPL/SFL & IPL/GCL | 9.30 | p<0.001 | 0.98 |
| MCL | 16.73 | 10.13 | 100 | MCL & EPL/SFL | 2.60 | 0.009 | 0.27 |
| IPL/GCL | 5.14 | 7.22 | 147 | MCL & IPL/GCL | 9.69 | p<0.001 | 1.45 |

**Table 2:** Comparing theta, gamma, and spiking power relative to the MCL. Significant differences were observed on either side of the MCL for all frequency bands evaluated. Significant differences were also observed between the MCL and the IPL/GCL for all frequency bands, but only spiking power showed a significant difference between the MCL and the EPL/SFL, suggesting power changes were steeper on the deeper side of the MCL. To test for significant changes in power, a Wilcoxon’s rank-sum test was used for the average power as calculated within 240 µm of either side of the MCL. If there were less than 240 µm of electrode array, available sites were used to calculate the mean. If there were no electrode arrays on one side of the MCL then no measure for that side of the electrode array was included in the rank-sum analysis, thus the number of electrode arrays used to calculate the mean across the layers for each power is indicated (n). Effect sizes were computed using Cohen’s d.

|  | Mean | Standard deviation | n | Combination | rank-sum score | p-value | cohen's d |
| --- | --- | --- | --- | --- | --- | --- | --- |
| Theta Power |  |  |  |  |  |  |  |
| EPL/SFL | 0.28 | 0.19 | 32 | EPL/SFL & IPL/GCL | 4.24 | <0.001 | 1.39 |
| MCL | 0.37 | 0.31 | 34 | MCL & EPL/SFL | 0.67 | 0.5006 | *ns |
| IPL/GCL | 0.62 | 0.29 | 31 | MCL & IPL/GCL | 3.26 | <0.001 | 0.84 |
| Gamma Power |  |  |  |  |  |  |  |
|  |  |  |  | 31 |  |  |  |
| EPL/SFL | 0.23 | 0.22 | 32 | EPL/SFL & IPL/GCL | 5.79 | <0.001 | 2.27 |
| MCL | 0.17 | 0.20 | 34 | MCL & EPL/SFL | 1.48 | 0.1384 | *ns |
| IPL/GCL | 0.66 | 0.15 | 31 | MCL & IPL/GCL | 6.17 | <0.001 | 2.74 |
| Spiking Power |  |  |  |  |  |  |  |
|  |  |  |  | 31 |  |  |  |
| EPL/SFL | 0.52 | 0.20 | 32 | EPL/SFL & IPL/GCL | 5.79 | <0.001 | 2.27 |
| MCL | 0.77 | 0.20 | 34 | MCL & EPL/SFL | 6.11 | <0.001 | 2.52 |
| IPL/GCL | 0.22 | 0.16 | 31 | MCL & IPL/GCL | 3.76 | <0.001 | 0.60 |

**Figure 3:**
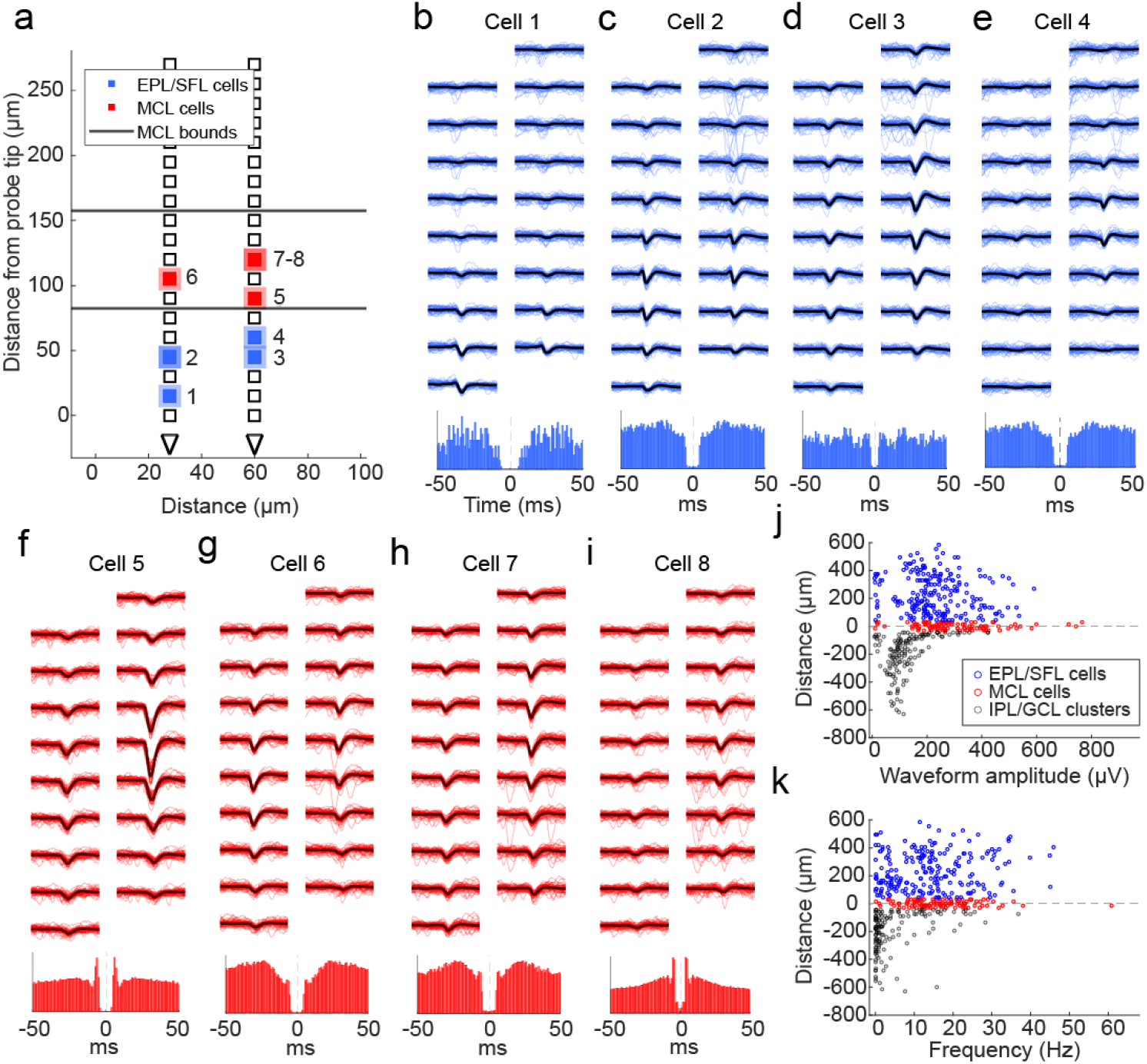
Physiological features of cells relative to the estimated MCL. **(A)** A schematic showing the location of MCL cells (red squares) and EPL/SFL cells (blue squares) across the recording sites of one example electrode array from a ventral OB recording. The edges of the estimated MCL are indicated by black lines. **(B)** (top) For a single EPL/SFL example cell, the mean waveform (black line) across 9 rows of recording sites (18 sites total) is plotted over 50 randomly selected individual waveforms (blue). The number above the plot indicates the cell’s location in (a). (bottom) The autocorrelogram of the cell is plotted (±50 ms) showing the cell displays a refractory period. **(C-E)** The same is plotted for three more EPL/SFL example cells and **(F-I)** four example MCL cells. **(J)** A scatter plot of each cell’s distance from the center of the MCL against its waveform amplitude from cells across all sessions shows amplitude declines moving toward the IPL/GCL, as displayed by the lower amplitude of IPL/GCL clusters (gray) as compared to MCL cells (red) and EPL/SFL cells (blue). **(K)** The same relationship can be observed for the average spiking rate of cells/clusters across the OB layers LFP (2-10 Hz) amplitude plotted across 500 ms from recording sites across 1 of the 4 probe shanks from a single recording session. Theta LFP, extracted using CWT based method, exhibits polarity reversal in the vicinity of the estimated MCL (red dashed line).

After defining layers, the spiking rates of sorted units were compare between layers. Spiking rates dropped moving away from MCL and deeper into the IPL/GCL, but spiking rates did not change to the same degree when moving in the direction of the superficial layers. The mean spiking frequency of all sorted units in the MCL (m = 16.73 Hz, std = 9.05) and in the EPL/SFL (m = 14.03 Hz, std =10.13) were significantly different from clusters in the IPL/GCL (m = 5.14 Hz, std = 7.33) as determined by rank-sum tests (Z = 9.69, p < 0.001, d = 1.45 and Z = 9.30, p < 0.001, d = 0.98 respectively). The MCL and EPL/IPL firing rates were significantly higher, but the effect size was small (Z = 2.60, p = 0.02, d = 0.27). Therefore, we found spiking rates as calculated across the entire recording session to be highest in the MCL and slightly lower in EPL/SFL.

It is to be noted that clusters found in the IPL/GCL are not multi-unit activity clusters, but rather seem to be good units from the IPL/GCL that exhibited physiologically feasible waveforms and refractory periods (Extended Data Fig 3). Despite this, they were not included in the analysis. In fact, it is likely that some clusters with high amplitude near the deep edge of the MCL (Fig 3j-k) are MCs mislabeled due to imperfect alignment or are cells with somas located in the IPL or superficial GCL such as deep short axon cells. Despite this, given the difficulty of differentiating between axonal activity and interneurons when sorting low amplitude waveforms from in vivo extracellular recordings (Barry, 2015), we refer to IPL/GCL units as clusters instead of cells and do not include them in further analysis.

### MCs and EPL/SFL cells show large-scale temporal response features

Despite unique trial to trial plume dynamics, responses averaged across trials within high and low concentration conditions showed a variety of response profiles to the larger temporal structure of the plume (Fig 4). Smoothed kernel density functions (KDFs) calculated in the style of Bolding and Franks (2017) were used to visualize cell response profiles, the cell firing rates across trials (Bolding & Franks, 2017). Some cells’ responses showed bursts of transient activity, while other cells exhibited longer and drifting responses across the plume (Fig 4b-c). The most dominant measurable features of trial-averaged responses seemed to be driven by macro-level temporal features of the plume, namely onset and offset. Onset responses and offset responses are defined as a significant change in binned spike counts relative to baseline firing rates using a Mann-Whitney U-test. Plume onset and offset were both manually scored. Plume onset was defined as the first 500 ms after the first whiff of the plume, and plume offset as defined as the first 500 ms after the last whiff of the plume ends. For offset responses, this means that a response that returns to baseline at offset will be considered non-significant, but responses that rebound after the plume and responses during the plume that do not terminate their responses at plume offset will both be considered offset responses. Thus, offset responses may signal the exact timing of odor offset by rebounding or may maintain a neural representation of the odor after offset by sustaining. Across all trials, we found 13.5% of cells showed onset responses and 7.6% of cells showed offset responses. Using a Wilcoxon signed-rank test, we found that the number of cells exhibiting plume onset and offset responses did not differ significantly across sessions (p>0.05).

**Figure 4:**
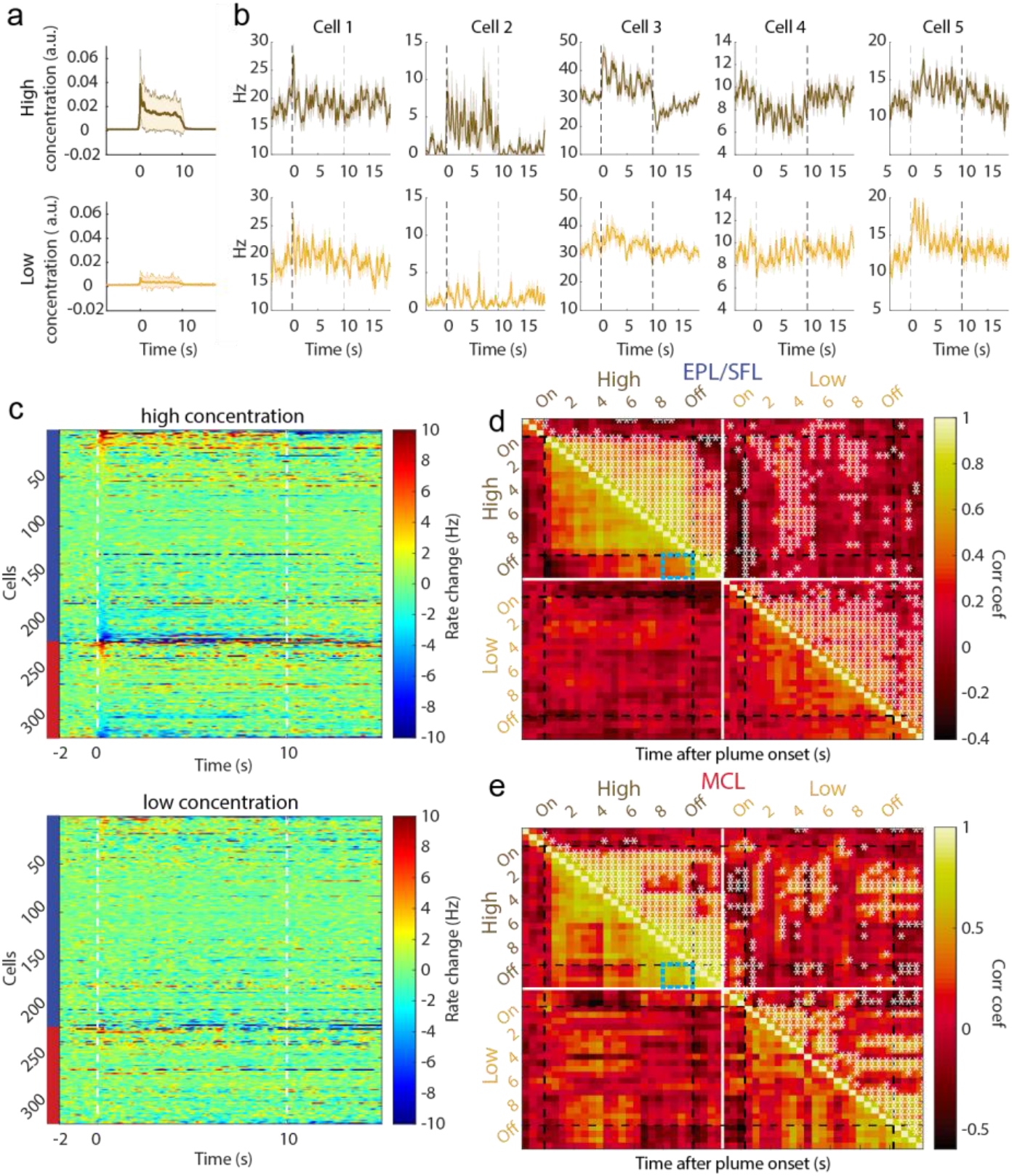
Cells exhibit different large-scale temporal response features to plume presentations such as characteristic onset and offset responses. **(A)** Odor across all trials is aligned to plume onset and the average odor concentration for each timepoint within low and high concentration trials is plotted (mn +/- 1std) showing the relative difference between low and high concentration. **(B)** The trial averaged firing rate of example cells are plotted (mn +/- 1std) showing that despite each trial having unique concentration dynamics, response features emerge on larger timescales that differ between cells. **(C)** The trial averaged responses are plotted as the change in firing rate from baseline (-2s to -1.5s) within high concentration trials (top) and within low concentration trials (bottom). (Color axis is cropped to 10Hz ensure cells with smaller firing rate changes can be visualized in the plot.) Cells are sorted by plume onset (‘on’, 0-500 ms) within EPL/SFL cells (indicated by left side blue bar) and within MCs (red bar). **(D)** The response of EPL/SFL cells (top) compared across time are plotted for high concentration trials to see how responses evolve across plume presentations. The trial-averaged firing rate for each cell is calculated for each 500 ms, resulting a single array with each cell’s mean rate for every time window. Cell activity is correlated across plumes for EPL/SFL cells and notably decorrelated from baseline responses. Significant correlations between time bins are indicated in the upper triangle with asterisks (p<0.05). Correlation coefficients showing the relationship between the end of plume response and offset response are noted for both cell types on high concentration trials (blue boxes). **(E)** Same as (d) but for MCs. MCs showed high correlations between their activity at the end of the plume and their activity after plume offset when odor is no longer present.

When comparing responses across concentration conditions, we found that responses were elicited more often in high concentration trials (onset 13.2%, offset 12.9%) than low concentration trials (onset 6.6%, offset 4.1%). The level of observed offset responses in low concentration trials (4.1%) suggests that plume offset responses are not significantly elicited by the lower concentration plumes as the number of responses observed did not exceed the expected level of Type I error (p<0.05 threshold).

Previous research on sustained responses has not differentiated between MCs and TCs, and our data suggest that despite similar responsivity levels, EPL/SFL cells may be less likely to show sustained responses and more likely to show rebound responses than MCs (Extended Data Fig 4 c-d). A cell response that sustains should show correlated activity between the end of the plume and after plume offset. Correlations between time windows (500 ms) across cell responses show differences between the two cell groups in how related offset responses are to end of plume cell responses (Fig 4d-e). When correlations are measured during high concentration trials that exhibit offset responses, correlation coefficients between the end of plume time bins and plume offset time bins are significantly different (Figure d-e, blue boxes). To quantify these differences between offset responses in EPL/SFLs and MCs, we calculated correlations using only EPL/SFL cells and MCs with significant offset responses as compared to baseline (Extended Data Fig 4). Correlation coefficients between the windows of spiking activity in the first two seconds after offset and windows from the last 2 seconds prior to offset were highly correlated for MCs (µ=.90, std=0.07). They were also significantly more correlated than they are for EPL/SFL cells (µ=.67, std=0.1) as determined by a Wilcoxon rank sum test (p<0.001, z=4.3908). Higher similarity between plume end and offset responses for MC cells suggests MC cells are more likely to sustain responses at plume offset and less likely to show rebound responses that EPL/SFL cells.

### Plume Responsivity

We measured responsivity to plume presentations at the cellular level using the ZETA-test (Fig 5, (Montijn et al., 2021)). As the stimulus dynamic did not repeat across trials, there is no reason to assume cell dynamics would repeat. With fewer a priori assumptions for alignment and averaging, choosing a meaningful z-score threshold and meaningful time bins across the 10 second plume becomes a more difficult problem. ZETA analysis exhibits increased sensitivity in the detection of significant responses without the need for optimizing parameters such as timescales for binning spike counts or for averaging when compared to other standard mean driven analyses. ZETA analysis can be conceptualized as a K–S test with a bootstrapped null confidence interval which allows one to determine if an observed cumulative distribution function (CDF) of a cell’s response significantly diverges from linearity (the CDF of a constant baseline firing rate is linear). Thus, if the spike times diverges enough from linearity, a cell can be classified as having a significant response. Therefore, we used the ZETA analysis to determine which cells had significant responses to naturalistic olfactory plumes.

**Figure 5:**
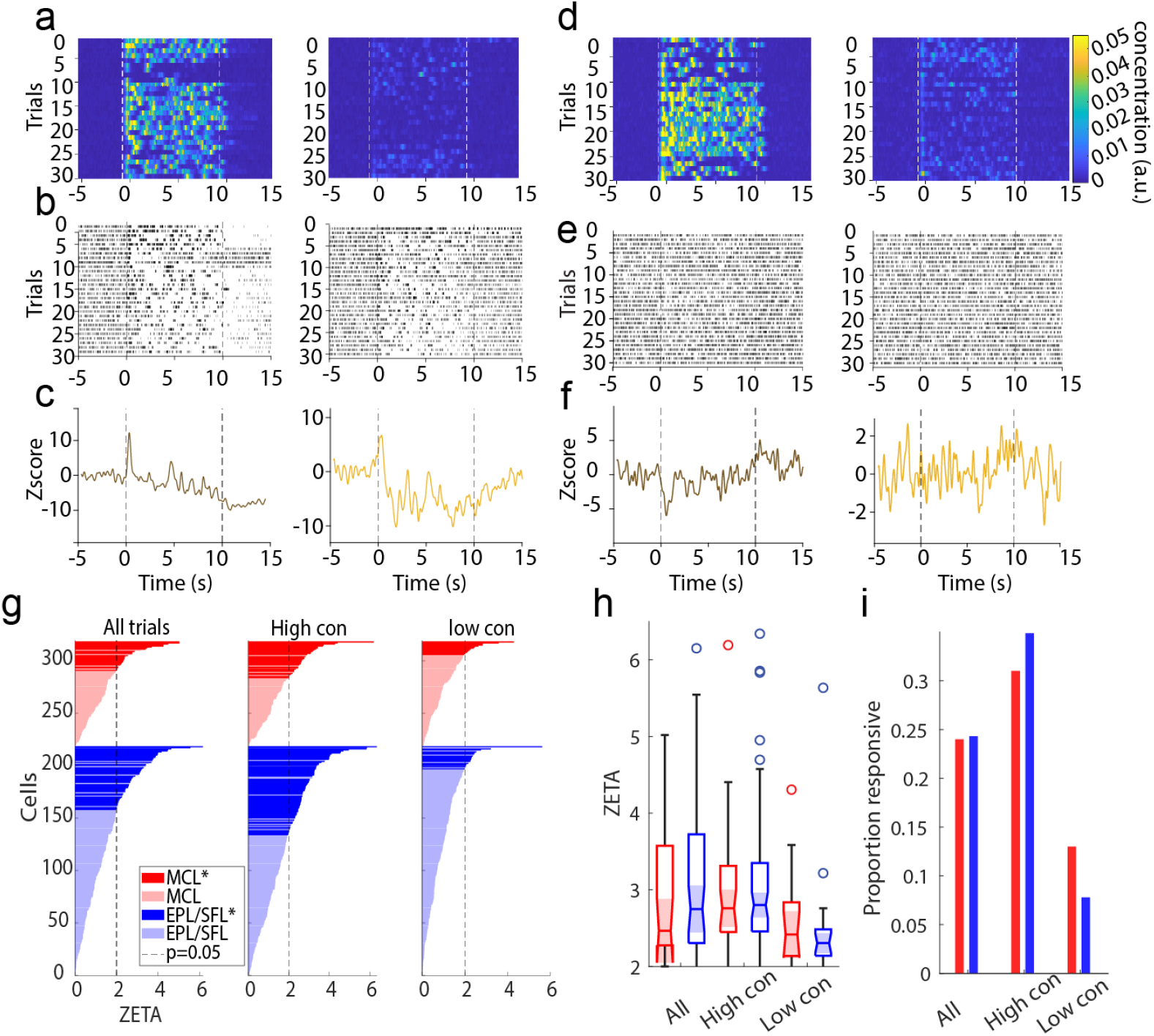
Significant responsivity to plume presentations as measured by ZETA analysis. **(A)** Odor concentration aligned to plume onset is plotted for all high concentration trials (left) and all low concentration trials (right). Onset (0s) and estimated offset (10s after onset) are indicated dashed white lines. **(B)** Raster plots and **(C)** PSTHs are plotted for a single example cell exhibiting a reliable, robust, and sustained response across high and low concentration trials and its response profile represents a small minority of the dataset. **(D-F)** The same is plotted for a cell that exhibits a response profile of lower magnitude and which is more representative of the majority of responsive cells in the dataset. **(G)** ZETA values are plotted for all MCs (red) and EPL/SFLs (blue) as calculated across all trials (left), within high concentration trials (middle), or within low concentration trials (right). **(H)** The distribution of ZETA scores is plotted for significantly responding cells as calculated within each condition and distributions do not significantly differ across conditions. **(I)** The plotted proportion of significantly responding cells for each condition shows responsivity does not significantly differ between cell types within any concentration condition. Higher concentration trials elicit more cell responses from the population.

To perform this analysis, spike times are translated into the spike’s latency from the plume onset time of its corresponding trial. The CDF is then calculated from the onset relative spike times from all trials. Cells were considered to respond significantly to plumes when their ZETA score indicated spiking activity significantly deviated from a linear baseline (see methods). Cells were considered to have a significant response only if, in addition to their significant ZETA value, the timing of either the positive or negative linear deflection maximum (ZETA score) occurred during the plume period (defined as lasting from plume onset until 1 second after plume offset, 0-11s). Therefore, cells with significant ZETA scores in the pre-plume(-5-0s) or post offset period (12-15s) were not considered responsive.

In the MCL, 41% (41/100) of cells significantly responded to plumes and in the EPL/SFL 43.12% (94/218) of cells significantly responded (Fig 5g). MCs and EPL/SFL cells show similar responsivity to plumes as neither the ZETA score magnitude (Fig 5h) nor the proportion of significantly responsive cells (Fig 5i) differed significantly when compared across all trials, across only high concentration trials, or across only low concentrations with sessions (Wilcoxon signed-rank tests p>0.05 for all comparisons). Thus, MCs and EPL/SFL cells showed similar plume responsivity.

### The majority of OB cells do not resolve odor concentration dynamics with high fidelity

Responsivity levels, as measured by ZETA analysis, do not account for stimulus properties aside from the timing of the plume onset, which is used to define their latency-defined spike times. Therefore, responsivity scores do not address whether cells follow odor concentration changes across time. To address this question, correlation analysis was used to quantify how well individual cells resolve plume dynamics (Fig 6a-b). For each cell, the correlation between a cell’s spike rate and the odor concentration across the plume were calculated for each trial (see methods). Correlation coefficients were then averaged across all trials, across only high concentration trials, and across only low concentration trials. For each cell, coefficients were compared to their respective 95% null confidence interval from a trial shuffled bootstrap analysis to determine if the correlation was significant. Across all cells, 105/318 cells had above chance level correlations with odor concentration on all trials (µ=0.06, std=0.03), 119/318 on high concentration trials (µ=0.09, std=0.04), and 32/318 on low concentration trials (µ=0.07, std=0. 03). Correlations were weak: only a minority of cells had coefficients of magnitude greater than 0.1 (Fig 6b). The number of cells with coefficients above 0.1 magnitude was 8/105 cells across all trials, 31/119 cells across high concentration trials, and 2/32 cells across low concentration trials. Neither the proportion of significantly correlated neurons (Fisher’s exact test, p>0.05) nor the magnitude of those correlations differed between MCs and EPL/SFL cells (Wilcoxon signed-rank test, p>0.05). This suggests that despite having decreased sensitivity to odor concentration (Burton & Urban, 2014) and less direct input from OSNs relative to TCs (Gire et al., 2012; Gire & Schoppa, 2009; Najac, de Saint Jan, et al., 2011), MCs exhibited comparable ability to follow odor concentration dynamics across a plume.

**Figure 6:**
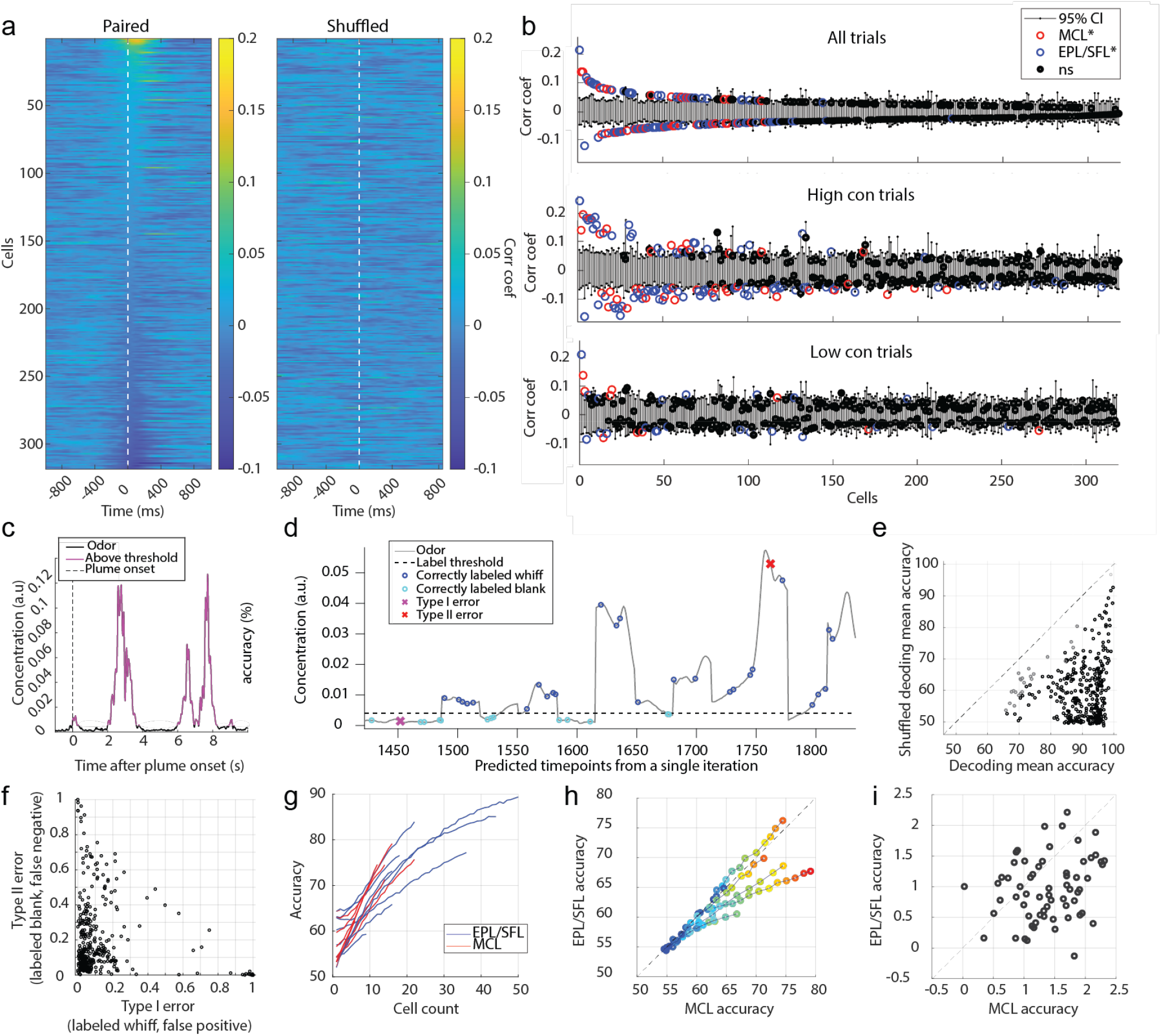
Despite weak correlation between individual cell responses and odor concentration dynamics, populations decode odor concentration with high accuracy. **(A)** The cross-correlation between the odor concentration and each cell’s spiking rate is calculated within each trial and then averaged across trials. Each row depicts the mean correlation coefficient between the cell’s spiking rate and the odor concentration for each indicated lag ±1 s. Cells are sorted in order of decreasing magnitude of the max mean correlation coefficient within 0-500 ms lag of neural activity following odor concentration. **(B)** The mean correlation coefficients of all cells are plotted against their respective bootstrapped 95% confidence interval for all trials (top), for high concentration trials only (middle), and for low concentration trials only (bottom) and 30.2%, 37.4% and 10.1% (respectively) of cells were weakly but significantly correlated. **(C)** A binarized odor signal was used to test decoding accuracy of OB spiking activity at the population level to discern whiffs (above threshold) and blanks (below threshold). A single example trial is plotted showing the odor concentration of the plume binarized as blanks (black) or whiffs (magenta). **(D)** Predicted labels are plotted over the odor concentration of timepoints from an example classifier from a middle of the plume time window (5s -5.25 s). Nonlabelled samples were used as training data. **(E)** The mean decoding accuracy for radial kernel SVM classifiers run across the 40 time bins (250 ms binned windows from onset to offset) from each session show the majority of cell populations decode odor presence at above-chance level accuracy (black) across the duration of the plume. Grey points indicate time windows for which the mean classifier accuracy did not exceed the mean accuracy its respective shuffled confidence interval. **(F)** The mean Type I and Type II error rate is plotted for each classifier suggesting that classifiers tend to bias the errors towards misses and are more likely to incorrectly label a whiff as a blank than vice versa. **(G)** The mean decoding accuracy of radial kernel SVM classifiers across the entire plume are plotted for the MCL (red) and EPL/SFL (blue) populations for sessions and calculated across increasing sample sizes within each cell type. MCL populations exhibit sharper accuracy increases per added cell than EPL/SFL populations compared within session. **(H)** To directly compare changes of decoding accuracy of the two cells type within sessions using matched sample sizes, the mean accuracy of MC and EPL/SFL populations are plotted against each other for increasing samples sizes. The first dark blue circle indicates the smallest sample size of 2 neurons for sessions and each shade closer to red indicates an increase in the population size by 1 neuron. (For each sessions, the largest matched sample size is constrained by the minimum number of cells across the two types). **(I)** The mean gained accuracy for each added MC and each added EPL/SFL is calculated across all sessions (calculated from the derivative of mean accuracy curves plotted in (h)). On average the addition of a single MC to a population added significantly more explanatory power than the addition of an EPL/SFL cell as exhibited by the majority of points falling below the diagonal line.

Although a subset of cells weakly tracked continuous concentration changes, cells may still encode odor presence versus absence with relatively higher fidelity. It is possible that cell responses that do not closely align with concentration may still respond differently during odor whiffs than during odorless periods. To examine this at the individual cellular level using correlation analysis, odor was thresholded into odor on and odor off periods (see methods, Fig 6c) and correlation coefficients were calculated between neural firing rates and the binarized odor signal across the plume (Extended Data Fig 5). The distribution of correlation coefficients for cells when calculated using the binary odor signal was not qualitatively different from coefficients calculated using the full odor concentration dynamics. Therefore, at the individual level, cells do track odor presence versus absence with high-fidelity in a concentration invariant manner. This indicates that, in general, individual cells may not respond as consistently to whiffs across a plume as they do to isolated odor pulses.

In summary, a portion of the population exhibited weak but significant correlations with plume dynamics, suggesting many cells in the OB respond to odor during plume presentations but do not reliably follow concentration fluctuations across plumes at the individual level.

### OB populations discern whiffs and blanks across the duration of a plume at the population level

Despite this lack of high-fidelity tracking of plume dynamics at the cellular level, we next tested whether cells resolved odor at the population level and found whiffs and blanks could be accurately discerned across the plume using a nonlinear decoding method. To test whether population activity was significantly different during periods with versus without odor, Support Vector Machine (SVM) binary classifiers were used to measure the accuracy with which these periods could be decoded from OB population activity. Linear decoding performed above chance level for some, but not the majority, of sessions (Extended Data Fig 6 a), but even those where linear decoding was above chance level, decoding was much more robust using a nonlinear kernel (Extended Data Fig 6 b-f). Odor concentration time series were binarized (Fig 6 c) and MCs and EPL/SFL cells together were trained to classify timepoints as either odor on (above threshold) or odor off (below threshold). To determine chance level decoding, decoding accuracy was compared to a shuffled 95% confidence interval.

The majority of classifiers discerned whiff and blank timepoints significantly above chance level (Fig 6d-e), with 334/360 time bins across sessions having mean decoding accuracy exceeding a shuffled confidence interval (Extended Data Fig 6, see methods). Accuracy was high (µ=89.73%±7.48% std), with performance exceeding chance level by ∼30% on average across sessions (µ=29.85%±10.99% std). Many cells exhibit drift or a wane in their mean response between plume onset and offset (Fig 4b-c), therefore, decoding accuracy was measured for each 250 ms window from plume onset (0s) to plume offset (10s). Despite changes in mean cell responses as a function of plume duration, population level decoding accuracy did not decrease as a function of plume duration and stayed robustly above change detection across the plume (Extended Data Fig 6). In general, classifiers were more likely to make Type II error (Fig 6 f), meaning that whiffs were more likely to be classified as blanks than vice versa. The same was error bias was found for linear classifiers (Extended Data Fig 6c). Our results suggest that although individual cells do not track concentration well, odor presence, whiffs, versus absence, blanks, could be decoded accurately from the population activity.

To determine the relative contribution of MCs and EPL/SFL cells to decoding accuracy, accuracy was assessed within cell type across populations of increasing cell count (Fig 6g). This determined how decoding changed as a function of cell count within each of the two groups (see methods). To quantify the average contribution of each added cell to the population decoding accuracy, we took the average change in mean performance with each increase in population size for both cell types. The mean rate of accuracy increase per cell was larger for MCs as compared to EPL/SFL cells as measured by a K-S test (*D*=0.45, *p*<0.001). MC populations were typically smaller than EFL/SFL populations for recording sessions. The addition of a cell to a small population will likely yield a higher increase in performance than the addition of a cell to a larger population. To ensure that the smaller size of MCL populations did not drive the observed results of MCs having higher mean contributions to decoding accuracy, an additional analysis was performed where EPL/SFL curves were truncated at the same size as their corresponding MCL population within each session. For the matched sample sizes, the additional accruacy with each MC was plotted against the additional accruacy for each added EPL/SFL cell for each sessions (Fig 6 h). The average change in mean performance with each added MC was plotted against the change in performance for each added EPL/SFL cell for the matched sample sizes (Figure 6 i). Results were qualitatively similar, showing that the addition of a single MC increases decoding accuracy more than the addition of a single EPL/SFLs when CDFs were compared using a 2-sample KS test (D=0.32, p<0.001).

As the decoded odor is binary and the threshold is set to discriminate when odor is present or absent during the plume, these results do not speak to differences in overall excitability between OB cells types. Tufted cells show higher excitability at lower concentration ranges (Burton & Urban, 2014; Chae et al., 2020). Our results speak to a possible difference between OB cells in tracking odor presence across intermittent plumes. Individual MCs may be more informative than individual EPL/SFL cells in discerning whiffs and blanks across plumes. It is clear though, that both cell types distinguish odor presence (whiffs) from odor absence (blanks), and properties of whiffs and blanks are tools that can be used to estimate odor source location (Pannunzi & Nowotny, 2019; Park et al., 2016; Rigolli et al., 2022; Yee et al., 1995).

## Discussion

### Summary

Using Neuropixels 2.0 probes to collect large-scale OB population activity and a modified odor sensor to assess real time plume dynamics, we have shown how natural plumes are encoded in the olfactory system of awake mice, examining both the cellular and population levels. Our work is the first to record plume dynamics at the nose while simultaneously capturing OB activity using high-density electrophysiology. In summary, cells respond to key temporal features of plumes, including plume onset, plume offset, and whiff-and-blank structure.

Our findings suggest individual cells signal when animals enter and exit plumes as they navigate their environment, but only weakly follow fine fluctuations in odor concentration. Despite an observed lack of high-fidelity tracking at the cellular level, SVM decoders demonstrated that cell populations accurately distinguish whiffs from blanks throughout the entire plume duration. Together, these findings at the cellular and population level support the idea that as a mouse navigates a plume, OB output relays macrolevel temporal plume features to higher olfactory cortex.

### Broadly encoded, large-scale temporal plume dynamics may support switching between behavioral motifs during olfactory search

Salient events, such as entering or exiting a plume, are foundational aspects of odor searches capable of triggering behavioral changes in navigating animals (Vickers, 2000). Our data indicate that while we did not observe fine-scale temporal tracking, large-scale temporal features are prominent in the OB activity elicited by plumes (Figs 4,5 and 6). It is known that MTCs show odor on and off responses to square odor pulses, but we did not observe individual cells with reliable odor off and on responses across a plume, as individual cells were only weakly correlated with odor across plumes. Instead, we found that plume onset and offset are globally encoded across the OB. Around 1 in every 10 cells had significant onset responses across all plume trials, and 1 in every 10 cells show offset responses when odor concentration is sufficiently high. This suggests that the OB broadly encodes the moment when an animal enters and exits a plume at the level of individual cells.

While mouse behavioral motifs during odor tracking are not as well characterized as those of invertebrates such as moths (Vickers, 2000), these global signals predict that similar behavioral switching could occur in mice upon plume entrance and exit during odor tracking. This idea that plume dynamics could cue switches in search behavior motifs is in good accordance with recent computational work investigating behavioral switching during simulated airborne-odor tracking (Rigolli, Reddy, et al., 2022) and recent work showing changes in behavior result from plume encounters (Tariq et al., 2023)

Onset and offset responses were elicited by high concentration plume trials, but not low concentration trials. The increase in onset responses during high concentration trials is consistent with the idea that higher concentrations recruit stronger glomerular responses. The increase is offset responses for high concentration conditions is consistent with previous research that MTC responses which persist after odor offset grow stronger as odor concentration increases (Patterson et al., 2013). As odor concentration increases with proximity to an odor source (Webster & Weissburg, 2001), our data suggests that plume onset and offset responses increase as an animal nears odor sources. Mice are more likely to switch to odor-guided search as they grow nearer to odor sources, and less likely to rely solely on other means of navigation such as memory or visual cues (Jackson et al., 2020). Increased OB activity regarding odor onset and offset near odor sources could be one of the tools that facilitates this behavioral switch.

It should be noted that odor offset responses also increase with the length of an odor stimulus (Patterson et al., 2013). In our study, concentration and intermittency (here defined as the proportion of time odor is above threshold) cannot be disentangled. High concentration trials were created using lower windspeeds. Lower windspeeds increase intermittency. Therefore, high concentration plumes also had higher intermittency, and odor was above threshold more often in these trials. Future research quantifying how concentration and intermittency each moderate plume offset responses is needed. This would inform when plume offset responses are strongest, and, in turn, how they may be used by olfactory navigators. Odor offset responses of MTCs have been previously documented in two ways. The first and more commonly reported odor offset response is that some MTCs exhibit rebound responses (Balu & Strowbridge, 2007; Cavarretta et al., 2018). We observed these classic rebound responses at plume offset (Fig 4b). A second and less well documented odor off response is sustained odor responses, referred to previously as olfactory ‘after-images’ (Patterson et al., 2013). These rebound responses are defined as sustained odor responses after odor is no longer present. We also observed sustained responses that continued in the same direction of the original odor elicited response, whether excitatory or inhibitory.

Plume offset responses observed were not all transient neural events, similar to previously reported. Longer odor offset responses to constant odor pulses have been observed (Patterson et al., 2013), and we found, that in addition to transient plume offset responses, some sustained offset responses were also observed. For EPL/SFLs and MCs with significant offset responses, we found significant correlations between end of plume responses and offset responses that extend for seconds after plume offset. This suggests plume offset responses are on a timescale consistent with short-term memory. Previous research has found that MTCs have odor offset responses after odor pulses. These responses can be used to decode odor identity and they scale with odor concentration (Patterson et al., 2013). The accuracy of odor representations diminishes over time following the odor pulse offset. This means that the OB may maintain representations of targets odor as animals navigate plumes even while navigating through blanks within a plume when odor is not detected. Additionally, this representation may wane as the time since the last whiff encounter lengthens. If so, waned plume offset responses could indicate that it has been too long since the last whiff was encountered, and a switch in behavior or search strategy is necessary. From a navigating mouse’s perspective, a period without odor could mean the animal is experiencing a blank within a plume or could mean the animal has exited the plume and is out of the range of further whiffs. The longer the blank, the more likely the second scenario becomes and the more likely a change in search behavior is needed, such as an aggressive turn back towards the lost plume. It is possible the number of cells still exhibiting sustained responses and the strength of those sustained responses may help discern the difference between a blank within a plume and an out of range positioning. Previous odor offset research did not specify between MCs and TCs, and our findings show that the sustained relationship of these offset responses was stronger for MCs. Thus, MCs may play a special role in maintaining target odor representations even when a mouse loses the air-borne odor trail.

Sustained and rebounding offset responses may also work together to provide important cues for tracking behavior. Rebound responses could signal the exact timing of when the last whiff encounter ended, and sustained responses, as mentioned, may help keep a neural correlate for time since the last whiff encounter in the OB. Hence, these two types of offset responses may both signify the disappearance of the target odor while retaining a form of short-term memory of the target odor as mice navigate plume-guided searches. This suggests that offset responses may play an informative role in tracking plume edges as mice navigate closer to the odor source and the plume cone narrows.

### The OB encodes whiffs and blanks across plumes

In turbulent plumes, “whiffs”, or periods of odor contact are interspersed with “blanks”, or time in which animals do not encounter odor (Celani et al., 2014; Riffell et al., 2008). Blank and whiff duration is known to vary with distance from an odor source (Celani et al., 2014; Murlis et al., 1992; Young et al., 2020). As one traverses downwind away from the source, whiffs begin to spread apart (Murlis et al., 1992) and each whiff expands (Young et al., 2020) resulting in a more diluted plume. Put another way, both blank and whiff duration increase, and there is a relationship between blank and whiff duration and distance from the odor source. Thus, the temporal structure of these encounters is a useful signal for source localization (Vickers, 2000).

We show that whiffs and blanks are reliably decoded across populations of MTCs, even when the firing of individual cells demonstrate weak correlations with the odor plume (Fig 6). Interestingly, our data suggests MCs and EPL/SFL cells may differ in the amount of information they carry regarding the presence of odor during a plume, with individual MCs carrying more information than individual EPL/SFL cells (Fig 6f). Since these cell types target different arrays of downstream structures (Igarashi et al., 2012), with TCs restricted to anterior piriform cortex (APC) and MCs projecting to a diverse collection of cortical and subcortical areas, including across the piriform cortex (PIR) and to the lateral entorhinal cortex (LEC), the difference in plume information content per cell has functional implications. As results are predominantly driven by a subgroup of sessions, future studies exploring the degree to which these cell types differ in the ability to resolve odor across plumes and its implication for downstream targets would be useful.

### A possible role for piriform cortex in odor-guided navigation

Convergence of multiple MTCs onto individual pyramidal cells in the anterior olfactory nucleus (AON) (Brunjes et al., 2005) and the PIR (Franks et al., 2011) could broadly convey whiffs and blanks to downstream targets. The rate of convergence of MTCs onto AON pyramidal cells have yet to be quantified, but the rate of convergence in the PIR has been estimated to be 200:1. This number is an order of magnitude higher than the number of cells we observed to be necessary for accurate decoding. Populations of only around ∼20 MTCs sufficiently discerned whiffs and blanks across the plume at above 90% accuracy. Since a radial kernel SVM decoder was required to achieve high decoding performance, the absence of robust linear decoding requires further investigation to determine how downstream targets may encode or interpret this signal. If PIR cells resolve whiff and blank timing from OB output, this opens the possibility that PIR carries a global whiff and blank signal for foraging animals.

It has been well documented that odor concentration, in addition to odor identity, can be decoded from PIR (Bolding & Franks, 2017; Stettler & Axel, 2009). Recently, a study showed how to infer the distance from a distant odor source using intensity and timing (including whiff and blank duration) dependent measures (Rigolli et al., 2022) and assessed the ability of these different cues to accurately predict odor source location. An important finding was that the combination of intensity and timing cues was more effective in predicting odor source location than the use of either cue type in isolation. Thus, it is possible that PIR integrates timing and intensity cues, which would be informative for localizing odor sources.

Lastly, odor place cells have recently been found in posterior PIR (Poo et al., 2022). The possibility of integrating odor identity, location, and plume cues (blanks, whiffs, and intensity) makes PIR a convincing target for future research into the neural mechanisms underlying olfactory search.

### Limitations for determining the limits of cellular level resolution of concentration dynamics

In this study, only a few cells were observed that exhibited correlation coefficients with comparable magnitude of those observed at the glomerular level in similar recordings using wide field imaging of glomerular activity (Lewis et al., 2021). This observation does not exclude the existence of cells that tightly track odor concentration, but correlations at the glomerular level observed using wide-field imaging techniques were greater than those observed at the cellular level using electrophysiological techniques.

It is possible that, aside from a more minor population of high performers, the majority of individual OB cells carry limited or nonlinear information regarding odor concentration changes or that OB cells respond to latent features of a plume that are yet to be tested.

One interpretation of these differences in tracking is that collective activity within a glomerulus is able to resolve concentration dynamics better than individual cells. This suggests that the average fluorescence of MTCs within a glomerulus has a stronger correlation with concentration dynamics compared to the activity of individual cells. Since we do not assign cells to glomeruli in our electrophysiological recordings, this question is outside the scope of this study as without identified sister cells from the same glomerulus we cannot assess the relative tracking ability of cells within a glomerulus. Another possible interpretation is that the widefield technique of Lewis et al., 2021 recorded from larger cellular populations across the bulbar surface, making it more likely to capture less prevalent, but highly responsive, glomeruli and that electrophysiological recordings without a targeted approach are less likely to capture this activity. Therefore, we cannot ascertain whether individual cells exist that track concentration to the same degree as observed at the glomerular level. However, we do note that our recordings did not sample any such cells.

Additionally, on a more technical note, the odor mixture used consisted of 5 odors known to show strong dorsal expression, but the majority of our recordings (8 of 10 sessions) were in ventral OB. Research on ventral expression levels of glomeruli is less extensive than that of dorsal or lateral OB areas due to the challenges associated with imaging ventral brain regions. The majority of our recordings were performed in ventral OB, as often spiking was not as strong in dorsal areas. This is in line with previous findings of stronger spiking in ventral OB (Paseltiner et al., 2020). Therefore, there may have been less highly responsive glomeruli in the ventral OB, leading us to capture fewer highly responsive MTCs. Regardless, information regarding large-scale features of the plume was still broadly encoded.

Our findings support the idea that the majority of cells in the OB do not resolve plume dynamics to the degree observed at the glomerular level, but does not speak to the limits of cellular level encoding of plume dynamics. Future studies could avoid this limitation by testing an odor panel before the session to target highly responsive cells or by using widefield imaging to target highly responsive glomeruli as the probe insertion site for proceeding electrophysiological recordings. Nonetheless, given that the number of highly sensitive glomeruli responding to each odorant is typically scarce, this study offers valuable insights into the responses of other constituent cells to natural olfactory scenes.

### Technical considerations of studied natural olfactory scenes and future directions

Plumes are stochastic due to turbulence (Celani et al., 2014; Riffell et al., 2008). This creates a significant problem for administering controlled and repeatable cues for the olfactory system.

In these recordings, we monitored input to the olfactory system using passive ethanol sensors. There are two things to consider given the accuracy of MOX sensors at measuring exact odor concentration levels at the mouse’s nose. First, sensors are placed with 4mm of the mouse’s nose and at that scale of distance (<5mm) there is some decorrelation of odor between any two spots in a plume (Tariq et al., 2021). Second, MOX sensors are robustly but not perfectly correlated with odor concentration as measured by photoionization detectors. The odor concentration at the mouse’s nose is estimated to be strongly correlated (∼0.6 Pearson correlation), but not perfectly correlated, with the signal measured by the sensor (Lewis et al., 2021). Therefore, the correlations and decoding accuracy reported in our findings should be considered to provide a lower bound estimate of the relationship between odor concentration across a plume and OB activity.

Since odor plumes are stochastic, difficulties exist in adapting statistical methods that are usually applied to repeated stimuli in the analysis of odor plume features. In general, we found that aligning to reproducible plume features was difficult, since fluctuations in odor concentration during a plume did not recur in the same context (due to the stochastic nature of the plumes) across multiple presentations. Olfactometers deliver repeatable odor stimuli, and recent development of olfactometers using high speed solenoid valves (Ackels et al., 2021) provides a way to design an odor concentration time series that mimics a stochastic plume and present this stimulus in a repeated and controlled manner. Changing air dilution over time using mass flow controllers to vary concentration, while also flickering fast solenoid valves final valves for fast transitions between whiffs and blanks, one may be able largely recreate the time series of odor concentration from a previously recorded plume.

Using reproducible stimuli with the full complexity of natural odor scenes is important. A problem with studying plume features exclusively in isolation is that recent stimulus history plays an important role in the processing of natural scenes. Presenting a stimulus embedded in dynamics relevant to natural odor environments is important, as it could help disentangle the effects of recent stimulus history across plumes. Odor representations are moderated by recent stimulus history as they can change over prolonged odor presentations or between sniffs even when a mouse smells a constant, unchanged stimulus (Baker et al., 2019; Fukunaga et al., 2012; Patterson et al., 2013; Spors & Grinvald, 2002). And varied odor off OB responses may moderate future responses depending on the size and intensity of recent whiffs within a plume. Additionally, neural activity in the OB has been shown to directly encode recent stimulus history, as some MTCs directly encode direction changes of odor concentration (Parabucki et al., 2019). Presenting dynamical features both embedded in plumes as well as isolated in dynamic motifs could help determine the degree to which encoding in the OB changes as a function of the recent history of plume dynamics. Just as other sensory systems have been documented to have more complicated encoding of stimuli in natural scenes than when those same stimuli are presented in isolation, it is important to understand how olfactory processing rules may change when odor is embedded in the complex dynamics of natural odor environments.

In addition to employing reproducible odor dynamics, an alternative approach involves harnessing optogenetics to stimulate cells in a time course that aligns with various aspects of plume dynamics. Temporally patterned optogenetic stimulus of the OSN terminals and simultaneous electrophysiology recordings of OB activity could help determine MTC responses to plume features in a reproducible and temporally precise manner. Although the advancement of optogenetic stimulus delivery methods (Chong et al., 2020) allows for this approach, the morphological complexity of the olfactory turbinates, as well as the specifics of receptor binding and signal transduction in OSNs are formidable obstacles to optically recreating naturalistic plume encounters. Additionally, divorcing sensory cues of the plumes from odor processing could affect responses. Odors are carried by fluid flow, which may create an inextricable link between plumes sensed by the olfactory system and local fluid dynamics, as sensed by the whiskers (Yu et al., 2016). Despite these caveats, optogenetics provide a promising avenue for future research of dynamic odor stimulus processing.

In summary, this work provides a pioneering attempt to understand some of the natural processing dynamics of the first olfactory relay in response to plume dynamics and considerations for future research.

## Acknowledgements

We would like to thank Anna Bowen for advice on Neuropixels data acquisition (Postdoctoral Fellow, Department of Biological Structures, University of Washington), and the Franks Lab (Neurobiology Department, Duke University) and Kelly Chang (Postdoctoral Fellow, Psychology Department, University of Washington) for consultation regarding data analysis. We would also like to thank Jesse Miller (Former Gire Lab member) for consulting on LFP analysis and Mohammad Tariq (Postdoctoral Fellow, University of Regensburg) for help with odor sensor setup. This paper was typeset with the bioRxiv word template by @Chrelli: www.github.com/chrelli/bioRxiv-word-template.

## Author contributions

DG and SL initiated the study and designed the experiments. SL and NS designed the rig and data acquisition pipeline for Neuropixels recordings. SL and LS performed the surgeries and experiments. DG, SL, NR, KF, and NS performed data analysis. SL and LS generated figures. SL, DG, NR, and NS wrote the manuscript.

## Declaration of generative AI and AI-assisted technologies in the writing process

During the preparation of this work the authors used ChatGPT in order to check for spelling and grammatical errors. After using this tool/service, the authors reviewed and edited the content as needed and take full responsibility for the content of the publication.

## Materials and Methods

### Mice

Experiments were performed on seven B6 mice (one female and six male) from the Jackson Laboratory (Strain # 000664) between 14-17 months of age. Animals were maintained on a 12-hour reverse light/dark cycle. All experimental procedures were approved by the Institutional Animal Care and Use Committee at the University of Washington.

### Olfactory Stimuli

Olfactory stimuli were released in a manner similar to Lewis et al., 2021 (Lewis et al., 2021). An automated odor port released odor within a 40cm x 40cm x 80cm acrylic wind tunnel where airspeed was controlled by a vacuum at the rear of the wind tunnel, posterior to the animal’s location (Fig 1a). Concentration dynamics of olfactory stimuli varied stochastically from trial to trial creating plumes with unique concentration dynamics on each trial. Plumes were created using the methods as described in Lewis et al., 2021, such that odor vapor was released upwind of the animal in the wind tunnel, and then a vacuum exhaust at the rear of the wind tunnel pulled the odor plume past the animal. To capture a variety of plume dynamics, flow was changed between low and high airflow speed in 10 block trials which resulted in trials with significantly different odor concentration levels. The location of the upstream odor port varied between either 13.5 cm to 18.5 cm upwind, and either 0 cm or 2 cm off centerline, but are not used as a factor in the analysis of neural responses. Each session consisted of 60 trials (except two sessions, 22 and 31 trials in duration respectively, were terminated early due to odor sensor acquisition errors). In each trial, mice passively experienced a 10-second-long plume presentation of the odorant mixture.

Ethanol concentration throughout each trial was measured in a similar manner to that described in Lewis et al., 2021 by using a modified, commercially available ethanol sensor placed within 3.5-4mm from the mouse’s right nostril (Fig 1b). Sensor signal was acquired at 125 Hz. The Short-Time Fourier transform of the ethanol sensor recording was taken using the Matlab stft() function and slow drift of sensor baseline was removed by setting all low frequency components under 3Hz to 0 before inverting the signal back using the Matlab istft() command. For all sessions, an odorant mixture of 5 odorants was used to try to increase the number of responsive glomeruli and ethanol was used as a tracer for the ethanol sensor to capture the odor concentration. The same mixture ratio (0.3% ethyl tiglate, 0.3% methyl tiglate, 0.3% allyl butyrate, 0.3% isobutyl propionate, 0.3% ethyl valerate, 92.8% 200 proof ethanol, and 5.7% distilled water) was used for 9 of the 10 sessions. For the first session of the 10 sessions a mixture with a slightly lower ethanol percentage was used (0.6% ethyl tiglate, 0.6% methyl tiglate, 0.6% allyl butyrate, 0.6% isobutyl propionate, 0.6% ethyl valerate, 85.7% 200 proof ethanol, and 11.3% distilled water), but the ethanol was increased for future sessions to optimize the odor concentration signal of the ethanol sensor. Odor solutions were stored in odor reservoirs (centrifuge tubes) with air-tight, customized tops. Tops had two openings connected by tubing. One tube was connected to a Clippard electric valve (part no. EV-2-12) to create airflow and the other was attached to a 3D printed odor port. For each plume presentation, the valve was opened to allow airflow into the tube such that odor vapors exited the odor reservoir and traveled through cylindrical tubing (1/16” inner diameter) to the release point at the odor port.

### Surgery

The mice underwent two surgeries prior to electrophysiological recordings. During the time between surgeries, animals were conditioned to the head-fix setup. For the first surgery, a cannula and head plate were implanted. After a complete recovery was made from surgery, mice were conditioned to the head-fix setup. The second surgery, a craniotomy over a dorsal area of the main olfactory bulb, was then performed. For both surgeries, mice (n = 7) were anesthetized with isoflurane for surgery.

During the first surgery, a custom-built cannula was implanted in the style of (Findley et al., 2021) over one of the mouse’s nostrils, although data from cannulas was not used in this study. Next, a customized stainless-steel head plate was glued directly on the skull centered near lambda coordinates. Metabond was then added to cover all exposed skull. Once the animal recovered from this surgery (2-3 days), handling and subsequent conditioning to the head-fix setup began.

Once the animal was conditioned to the head-fix setup (1-2 weeks), the second surgery, a craniotomy, was performed where a 1mm x 1.5mm rectangle was removed above one of the two olfactory bulbs. First, a dental drill was used to remove the Metabond over one bulb. Then, a small well was drilled into the Metabond caudal to the bulb to hold more ringer’s solution during the grounding (this protected the grounding solution from evaporating in the wind tunnel during recordings). A gold-plated socket (Newark D Sub contact socket, 66504-9) was attached with super glue into the bath space for the grounding pin. The craniotomy was then performed and then Kwik-Cast (Silicone Elastomer) was applied to seal off the craniotomy. Afterwards a small wall was built using rings of superglue each cured with Zip Kicker (CA accelerator) immediately after application to create a pool (Fig 1b) to hold the ground solution. After, a layer of Krazy glue was applied to the outside of the wall and cured with the accelerant to make sure the pool was leakproof. Last, the pool was gently rinsed multiple times with sterile saline solution.

### Electrophysiological Recordings

Four shank Neuropixels 2.0 electrode arrays were used to record OB activity during plume presentations. Probes were mounted on dovetail caps (uMp-NP2-CAP) with Metabond. The caps were attached to extension rods (uMp-NPR-200) by Neuropixel adapter heads (uMp-NPH). The extension rods were held by a four-axis micromanipulator (uMp-4).

Across the 5,120 possible recording sites, up to 384 can be chosen for simultaneous data acquisition. For all recording sessions, the bottom most 96 sites from each of the four shanks (the bottom 48 rows on each shank) were selected. The four shanks span an area of ∼750µm wide and ∼720µm deep, allowing for multiple layers of the OB to be captured simultaneously. Probes were inserted into one olfactory bulb such that the four shanks ran in a rostral to caudal direction along the sagittal plane of the OB. Recordings targeted either the dorsal (3/10) or ventral (7/10) OB. Recording sessions favored the ventral OB as stronger spiking activity was observed there, as has been previously reported (Paseltiner et al., 2020), facilitating stronger signal to noise ratio for isolating units in spike sorting.

Recordings were either performed on the same day as the craniotomy (n = 8 of 10) or on the day after (n = 2 of 10). For recordings performed on the same day, the mouse was anesthetized using isoflurane for surgery performed in the morning, and then recordings were performed in the afternoon once the mouse had fully recovered. Before the recording began, the Kwik-cast was removed and the grounding pool was filled with Ringer’s solution. Recording were made using an external reference, and for grounding, a gold-plated pin (Newark D Sub contact pin, 66506-9) soldered to an Ag wire (A-M Systems, No. 787000) was inserted into the socket in the grounding bath (see Surgery) with the other end of the wire soldered to the probe. The probe was then advanced into the OB and allowed to settle for 15-25 minutes prior to the start of the recording. For the single shank recording of spontaneously evoked activity used in (Extended Data Fig 1), surgery and electrophysiological acquisition are done in the style of Steinmetz et al., 2021. Data from the single shank recording shows that the findings of LFP and spiking near the estimated MCL from four-shank recordings spanning ∼750 µm (a distance shorter than the dorsal/ventral axis across most of the OB) qualitatively reflects activity near the estimated MCLs of a single shank spanning ∼1.8mm. This recording was done during an odor-less, spontaneously evoked period of activity. As this recording did not use the same experimental setup described in the methods, spontaneously evoked activity is plotted, but data is not included further in any analysis reported in this paper.

### Data Analysis

Neural recordings were acquired using SpikeGLX (https://github.com/billkarsh/SpikeGLX).

The data was next automatically spike sorted using Kilosort3 (https://github.com/MouseLand/Kilosort), and then manually curated using phy2 (https://github.com/cortex-lab/phy).

Units with contaminated refractory periods, unstable firing rates across the session, or waveforms indicative of multi-unit activity were marked as ‘Multi-Unit Activity’, and their use is explicitly labeled as such whenever discussed in the paper.

Matlab 2021a was used to analyze data and plot figures. Code by Nick Steinmetz from the Spikes repository (Waveform statistics and plotting: https://github.com/cortex-lab/spikes) and the npy-matlab repository (https://github.com/kwikteam/npy-matlab) were used, as well as code by David Tingley from the Buzcode Repository (CCG plots: https://github.com/buzsakilab/buzcode). reported in this paper.

### Analysis of Frequency Bands Using a Continuous Wavelet Transform Based Method

To estimate power across frequency ranges, we used a continuous wavelet transform method (CWT) as described in Cartas-Rosado et al 2020 (Cartas-Rosado et al., 2020). Amplitude plots and power estimates for each frequency band were calculated using neural activity across 5 seconds of the inter-trial interval, a time in which no odor or other stimulus was presented. The signal was first down sampled from 30 kHz to 1 kHz, and then transformed into the time frequency domain using the CWT Matlab function (cwt). The amplitude of these frequency ranges over time were extracted using the inverse continuous wavelet transform Matlab function (icwt). Frequency ranges extracted were theta (2-10 Hz), gamma (30-100 Hz), and spiking (300+ Hz) frequency ranges. Due to limits of the down sampled data, 434 Hz was the upper limit of the 300+Hz spiking frequency. The power for each frequency band was defined as the square of the root mean square amplitude values. The activity from an 8 second long clip was transformed and inverted, but only the middle 5 seconds of the clip were used for the power estimation to ensure all CWT coefficients used were located outside the cone of influence, protecting against known edge effects of the CWT.

### Estimation of the Mitral Cell Layer

The position of the MCL(s) across each shank was estimated by eyemanually based on previous findings of LFP changes near the MCL in gamma (30-100 Hz) and theta (2-10 Hz) frequency ranges and expected changes in the unique spiking activity (measured as 300 +Hz) of granule cells (GCs) which dominate the population in deep bulbar cortex (Extended Data Fig 1). Thus, the row of recording sites estimated to lie along the MCL was manually scored using the 3 parameters of gamma amplitude, spiking amplitude, and theta amplitude. Gamma amplitude was the most important factor in determining the location of the MCL as dipoles in gamma oscillations have previously been used to estimate the location of MCL previously in OB recordings without histology (Fourcaud-Trocmé et al., 2014). The second most important measure was spiking power, or amplitude in frequency ranges that capture cell spiking (300+Hz). As cortex located deeper than the MCL is dominated by axon-less GCs, which are thought not to spike during spontaneous activity (Cang & Isaacson, 2003), sharp decreases in spiking power were used. Last, theta power was used a corroborating evidence. Theta dipoles are not as strongly associated with MCL, but rather expected to reverse between the glomerular layer (GL) and the granule cell layer (GCL), which are located to superficial to and deeper than the MCL accordingly. Therefore, a nearby theta dipole should be reflected as nearby changes in theta power. Thus, coincident changes in gamma power, spiking power, and theta power were used to estimate the MCL location along each recording shank.

The strong oscillatory activity of the OB has been shown to exhibit observable relationships between OB layers and LFP characteristics, particularly in the form of LFP polarity reversals of gamma and theta oscillations. Dipoles are a known byproduct of transitions between sources and sinks in extracellular recordings. The OB has dipoles that can be observed in the LFP at both theta (Hu et al., 2022; Kay, 2015) and gamma frequencies (Rall & Shepherd, 1968; Rojas-Líbano & Kay, 2008; Wróbel et al., 2020). As gamma and theta frequencies are thought to be generated by unique circuitry in the OB (Fukunaga et al., 2014), the sources and sinks associated with these oscillations are not interchangeable. In our study, we searched for transitions looked for gamma and theta dipoles, as indicated by areas of low amplitude in the LFP, to aid in MCL estimation.

For gamma oscillations, polarity reverses near the MCL, between the EPL and the IPL (Rojas-Líbano & Kay, 2008). In previous research, the point of gamma polarity reversal in LFP across the depth of the bulb has been used to locate the MCL in extracellular LFP recordings without histological identification of the MCL (Fourcaud-Trocmé et al., 2014). Gamma, is often hypothesized to be a result of reciprocal dendrodendritic interactions between mitral and granule cells (Lagier et al., 2004; Rojas-Líbano & Kay, 2008). In support of this hypothesis, optogenetic silencing of GCs was observed to significantly lowers the power of gamma oscillations in the OB, despite not significantly altering theta power (Fukunaga et al., 2014). The dendrodendritic hypothesis suggests that the excitation of an MC excites GCs to which it has dendrodendritic connections. The GCs then inhibit the MC in return, as well as other MCs to which they share dendrodendritic connections. This creates a negative feedback loop and expands the influence of the inhibition to larger MC populations via lateral inhibition. Over a window of time, the excitation of MCs combined with the local negative feedback loop leads to alternating phases of excitation and inhibition periods in the gamma frequency range (Rojas-Líbano & Kay, 2008). These gamma oscillations create sinks and sources in the extracellular LFP that reverse polarity across the MCL (Rojas-Líbano & Kay, 2008). When the MCs are excited, there is a net influx of current to their dendrites and to dendrites of GCs they have excited, resulting in a sink in the LFP superficial to the MCL. In turn, deep dendrites and granule somas have a net outflux of current, creating a source in the LFP beneath the MCL. Thus, the sinks and sources of gamma oscillations lead to a polarity reversal near the MCL. Calculating gamma power across time, the area near the center of the reversal will display low amplitude. As the MCL should display less gamma power than its surroundings, creating a local minimum, transition to low gamma power were predominantly used to estimate the position of the MCL across shanks.

For theta oscillations, polarity reverses across the principal cells, between the GL and the GCL (Kay, 2015). Recent in vitro recordings across an entire slice of the olfactory bulb, found theta frequency primarily in the GL and GCL, corroborating the hypothesis that theta power should be lower in the EPL/MCL/IPL area. As the EPL/SFL side of the estimated MCL on each shank is only measured within 240 µm of the edge of the MCL, it is likely to consist largely of the EPL as opposed to more superficial layers as the EPL is around ∼200 µm in width (Hamilton et al., 2005). Therefore, although changes in theta power are not exclusively positioned within the MCL, theta power should decrease moving from the GCL to more superficial cortex before increasing again to high theta power in the GL. As theta dipoles are not as closely associated with the MCL location as gamma dipoles, decreases in gamma power were prioritized and decreases in theta power near the MCL was used as supporting evidence. Therefore, reductions in gamma power at the MCL and reductions in theta power in the vicinity of the MCL were both used as localizers.

In addition to dipoles, the distinct properties of OB cell populations exhibit an observable relationship across OB layers. Central OB layers located deeper than the MCL, the internal plexiform layer (IPL) and the GCL, consist predominantly of axon-less granule cells (Nagayama et al., 2014). As GCs are axon-less cells and rely on reciprocal dendrodendritic connections, the spiking activity observed in the GCL is generally dendritic spiking which exhibits lower amplitude waveforms in the extracellular LFP than axonal spiking (Häusser et al., 2000). Additionally, spiking power was evaluated only during spontaneously evoked activity, and the majority of GCs are not observed to spike during spontaneously evoked activity (Cang & Isaacson, 2003). Therefore, we expect spiking power should drop when crossing the MCL into deeper tissue. In fact, in a Neuropixels 2.0 single shank recording that spanned the entire dorsal to ventral length of the OB, we observed a central, pronounced area of low spiking amplitude sandwiched between the dorsal and ventral OB regions. Lower spiking power in deep bulbar cortex implies higher spiking power to be observed when moving from deeper cortex into the MCL. Therefore, sharp increases in spiking power are expected at the transition between IPL/MCL. It should be noted that shanks were not aligned to a local maximum in spiking power, but rather to a drop off or decline in spiking power that was coincident with an increase in gamma power and a nearby change in theta power. For example, a maximum in spiking power at the MCL or a plateau in power across both the MCL and the more superficial areas would lead to the same final alignment. Thus, a sharp increase in spiking power as a result of differing cell populations across OB layers provided the second important class of evidence for MCL estimation.

Taken together, coincident transitions to low theta power, low gamma power, and high spiking power were used to pinpoint the recording sites within the MCL. Once the MCL was estimated, two rows above the MCL and two rows below the MCL (±30µm) were labeled as falling within the MCL. All recording site rows more superficial than the estimated MCL edge are classified as EPL/SFL sites. Those deeper are referred to as IPL/GCL sites. If a shank did not have a drop in gamma power coincident with a rise in spiking power, the shank was labeled as not crossing the MCL. Data from these shanks were not used in the analysis as location within the bulb could not be estimated. Also, any sites showing a strong reduction in all power frequencies were discarded as this indicates the recording sites are above the dorsal surface.

### Observing Post Alignment LFP Power Across the Estimated MCL

We evaluated the relationship between gamma, theta, and spiking power across and between shanks by analyzing the correlation of the three measures across recordings sites and by evaluating the mean power changes around the MCL as averaged across shanks after alignment. Power was normalized between 0-1 for each shank within each frequency band so that only relative changes within frequency bands across each shank were considered.

We found reliable relationships between gamma, theta, and spiking power across recording sites used for the analysis. Power was estimated for all three band for each recording site and sites from all shanks were concatenated in order to form an array for each frequency band. Pearson correlation was then used to assess the relation between the three frequency bands. Mean gamma power was positively correlated with mean theta power for each electrode sites (p<0.001, r(1113) = -0.48), confirming electrode sites with higher gamma power were more likely to have higher theta power as well. Additionally, spiking was significantly inversely correlated with both theta (p < 0.001, r(1113) = -0.37) and gamma (p < 0.001, r(1113) = -0.60), suggesting that transitions to high spiking power were more likely to be observed with coincident reductions in gamma and theta power. Thus, there exists a relationship in the way these three oscillatory frequencies changes across the depth of recording shanks that encourages the idea that reliable coincident features can be determined across the 3 frequency bands.

We also evaluated the average power changes across all shanks around the estimated MCLs. To do this, power for each side was calculated using all sites within 240µm of either edge of the MCL. For the shanks in which the MCL was positioned such that there were less than 240µm of recording sites before the edge of the shank, the available sites were to calculate the average. If the MCL was at the edge of the probe and no electrode sites were available on one side of the MCL, only the sites on the available side were included for that shank in the analysis. As a result, samples on either side of the MCL across shanks were not strictly paired. Results were visualized by plotting normalized theta, gamma, and spiking powers (Extended Data Fig 2) for each electrode site against the depth of the site relative to the MCL to visualize the resulting relationship of gamma, theta, and spiking power after alignment. Theta power shows more variation, consistent with polarity reversal of theta being less closely associated with the position of the MCL as discussed previously. To quantify coincident transitions to low theta power, low gamma power, and high spiking power around the estimated MCLs as averaged across all electrode arrays and Wilcoxon ranksum test was used to access differences in power after alignment (Table 1). Power was also calculated across theta, gamma, and spiking frequencies for one single shank recording of spontaneously evoked activity (Extended Data Fig 1). A single shank traversing the dorsal/ventral axis will cross the MCL twice, and the single shank recording accordingly shows only 2 areas on the shank consistent with our assumptions of LPF activity near the MCL. These two locations, the resulting MCL estimates, both exhibit increases in spiking power and drops in theta and gamma power indicating polarity reversals.

As this study uses no histology and MCL estimates may have some variability, we do not assign other layers of the OB other than the MCL. Therefore, we do not try to distinguish between cells types outside of the MCL. Therefore, cells more superficial to the MCL are referred to as external plexiform layer and superficial layer cells (EPL/SFL cells), but do not distinguish cell type further to assign putative tufted or external tufted cell types.

### Waveform Analysis of Cells

To calculate the location of cells and their waveform amplitudes, 1000 waveforms were randomly extracted using the Spikes Repository (https://github.com/cortex-lab/spikes) function getWaveForms. The mean waveform was then calculated for each recording site. The recording site with the largest amplitude change between the minimum and maximum amplitude value was used to define the location of the cell or cluster across the electrode array. Waveform amplitude was defined as the maximum amplitude change across all recording sites.

### Mean Responses

Unless otherwise specified, mean responses, or response profiles, are defined by smoothed kernel density functions in the style of Bolding and Franks 2017, and responses were aligned to plume onset. KDFs were smoothed with a guassian kernel with a 100 ms standard deviation (Bolding & Franks, 2017). KDFs were first normalized using the mean and standard deviation across the entire KDFs, and then were baseline normalized using the mean and standard deviation across all timepoints from -4.5s to -1s relative to plume onset. KDFs plotted in the paper are all calculated aligned to plume onset. Plume onset was manually scored as the beginning of the first whiff of each plume.

For the calculation of ‘offset’ responses plotted in Extended Data Figure 4c-d, we also calculated offset aligned KDFs. We first manually scored plume offset as the end of the last whiff of each plume. This allowed for more precise ‘off’ means as the unique offset time of each trial is different. For consistencty, each cell’s offset KDF was normalized using the mean and standard deviation from their onset aligned KDF.

### Plume Responsivity of Cellular Responses

To determine the responses profiles of individual cells to plume presentations, an open-source package ZETA (https://github.com/JorritMontijn/ZETA) determine if the cumulative density function (CDF) of spike latencies relative to stimulus onset is significantly different from a linear (i.e. baseline) spike rate by employing a Kolmogorov-Smirnov based approach (Montijn et al., 2021). This approach avoids any binning or averaging across trials, and thus has been shown to be a useful tool for examining neural responses to natural scenes where there is not a priori knowledge of the relationship between the stimulus and the expected response profile. When testing Neuropixels recordings of cell populations responding to natural movies, the ZETA test exhibited similar performance to running multiple ANOVAs at different timescales. Thus, without having to optimize the timescale of binning or averaging, the ZETA analysis was used to determine significant responsivity of cells to the complex odor concentration dynamics of natural olfactory scenes.

For this analysis, spike times for each cell were first translated into their latency from plume onset of their respective trial. Only spikes that fell between the time window of 5 seconds prior to plume onset to 5 seconds after estimated plume offset (-5:15 s) were used for the analysis. The CDF of all spike times for a cell is assumed to be a diagonal line with a slope equal to baseline firing rate if the cell has a constant firing rate that does not change after plume onset. A bootstrapped null confidence interval (CI) is created by jittering plume onset times (±2s) and recalculating spike times as latency from the jittered onset to create a null distribution of CDFs. If the CDF significantly deters from linearity, meaning that it sufficiently exceeds the null CI, then the cell is considered to significantly respond to the plume. The most extreme deterrence from linearity in the positive direction is called the ZETA (Zenith of Event-based Time-locked Anomalies) peak and represents the point at which spiking maximally exceeds baseline firing rate. The negative peak of deterrence is called the inverse ZETA peak and represents when the firing rate maximally subceeds baseline firing rates. Thus, the timing of the ZETA peaks is not equal to response onset, as onset could occur before the maximal deterrence from linearity is reached. But if the maximum deterrence of ZETA was not reached between plume onset and 1 second after offset, the ZETA response was not considered significant regardless of its strength.

### Measuring cellular responses to plume concentration dynamics

Cross correlation was used to quantify the relationship between cellular responses and plume dynamics in the method used to calculate correlation coefficients between glomerular responses and plume dynamics in Lewis et al., 2021. Briefly, for every cell, the cross-correlation between the odor concentration and the cell spike rate was calculated for each trial. First, the spiking activity of the cell was binned to match the sample timepoints of the odor concentration (125 Hz) and then convolved with a gaussian kernel with a standard deviation of 100 ms. The odor signal and cell spike rate were then both normalized and the Matlab xcorr() function was used to calculate the Pearson correlation coefficient of the two signals for every lag. Coefficients were then averaged at each time lag across all trials of the indicated condition (all trials, high concentration trials only, or low concentration trials only). The maximum absolute mean coefficient within a 500 ms lag of the cell spiking lagging behind the odor concentration defined the correlation between cell spiking and plume dynamics. To determine significance, coefficients were compared to a 95% confidence interval calculated using a trial shuffled bootstrap analysis with 1000 iterations.

### Decoding odor presence with nonlinear binary SVM classifiers

To determine the ability of cell populations to discern whiffs and blanks across plumes, decoding accuracy of the binarized odor signal was measured by training and testing Support Vector Machine (SVM) classifiers. Each timepoint was an instance for classification and each cell’s spiking rate was a feature vector. The open source Matlab package libsvm (https://github.com/cjlin1/libsvm) was used to implement SVMs (Chang & Lin, 2011).

To binarize the odor concentration signal into whiffs (odor on) and blanks (odor off), an odor threshold was calculated for each session. Odor threshold was determined by dividing the odor signal into baseline periods (-9:-2 s before plume onset) and plume periods (plume onset to 10 s after onset). Threshold was defined as the mean of the baseline period plus 0.25 standard deviation of the odor on period (Fig 6c).

The spiking rate of the cell was first binned to match the sample timepoints of the odor concentration (125 Hz) and then convolved with a gaussian kernel with a standard deviation of 100 ms.

The binary odor signal was decoded using radial kernel SVM binary classifiers with 5-fold cross-validation using a split data ratio of 90:10 for training and testing, respectively. Decoding accuracy was measured two ways, first as a function of time across the plume presentation for the entire cell population and second as a function of cell number within MCL and EPL/SFL populations. For both analyses, all MCs and EFL/SFL cells were used for classification regardless of whether their were significantly responsive to the plume as measured by the ZETA analysis. The one exception is that one of the ten sessions was excluded from the analysis as only 7 cells were recorded and no cells exhibited significant plume responses. It should be noted that inclusion of this session does not qualitatively change any findings as mean classifier perfomance across sessions with this session included still accurately classifies whiffs and blanks 27.21% above chance level (as opposed to %29.86 above chance when this session is excluded). Pseudo-population level decoding accuracy was not measured as cells can not be pooled across trials since concentration time series are stochastic and are reproduced across neither trials nor sessions.

Decoding accuracy was first measured for different time windows across the 10 second plume to see if populations could discern when odor was present and, if so, whether this ability changed across the duration of the plume. Decoding accuracy of combined MCL and EFL/SFL populations was measured using 100 iterations of SVM classifiers for each 250 ms time window stretching from plume onset (0 s) to estimated plume offset (10 s). The classifier was run for each 500 ms window from onset to offset resulting in 40 time windows for classification. For each window, instances consisted of all time points from all trials that fell within the designated time window. For example, for the first window (0-250 ms), all time points from the first 250 ms of the plume for each trial were used.

To determine if decoding accuracy was above chance level, bootstrapped confidence intervals were calculated. As the exact proportion of the time odor was above threshold not 50%, bootstrapped 95% confidence intervals of 100 iterations of 5-fold cross-validated classifiers defined chance level performance. For the shuffled analysis, binary odor labels of the training data set were shuffled. Odor presence was particularly skewed at the very beginning and end of the window due to the alignment process, therefore chance detection tended to be higher at these times. More specifically, for the 0-500 ms window, odor is often on more than 50% of the time due to plume onset being defined as the start of the first whiff of each plume. Since the estimated plume offset is set at 10 seconds after onset, odor tends to be absent when nearing estimated offset. This is because in a sparser plume, odor has begun to be released from the upstream odor port before a whiff comes into contact with the sensor. The odor port will always close 10 seconds after it first opens, not 10 seconds after the first whiff is measured at the sensors location. Therefore, the last second before estimated offset tends to have less odor than the middle of the plume. Thus, chance levels are assessed relative to the bootstrapped null confidence window for each window.

To determine the relative contribution of each MC cell to decoding accuracy as compared to the contribution of each EPL/SFL cell, decoding accuracy within each cell type was measured for each session. Classifiers were run for multiple population sizes ranging from a single cell to the full number of MCs or EPL/SFL cells recorded in the session. As decoding ability did not seem to vary as a function of plume duration, a single time window was used for classification which included all timepoints from onset to offset across all trials. For each possible cell sample size, 100 iterations of the 5-fold cross-validated SVMs were performed. This resulted in a mean decoding accuracy curve across iterations for both MCL and EPL/SFL populations within each session as a function of cell number (Fig 6g-h). For each session, the derivative of the mean performance curve was used to quantify the average change in decoding accuracy each time a cell was added to the sample size (Fig 6i). The derivatives for all sessions were then concatenated to create an MC distribution and an EPL/SFL distribution. In this way, the two distributions measured the mean increase in decoding performance achieved by adding one cell to the sample size (Gig 6f). The mean rate of accuracy increase per cell was then compared between MCs and EFL/SFL cells to assess the relative contribution to classifier accuracy of adding an MC as compared to adding an EFL/SFL cell.

## Extended Data Figures

**Extended Data Figure 1:**
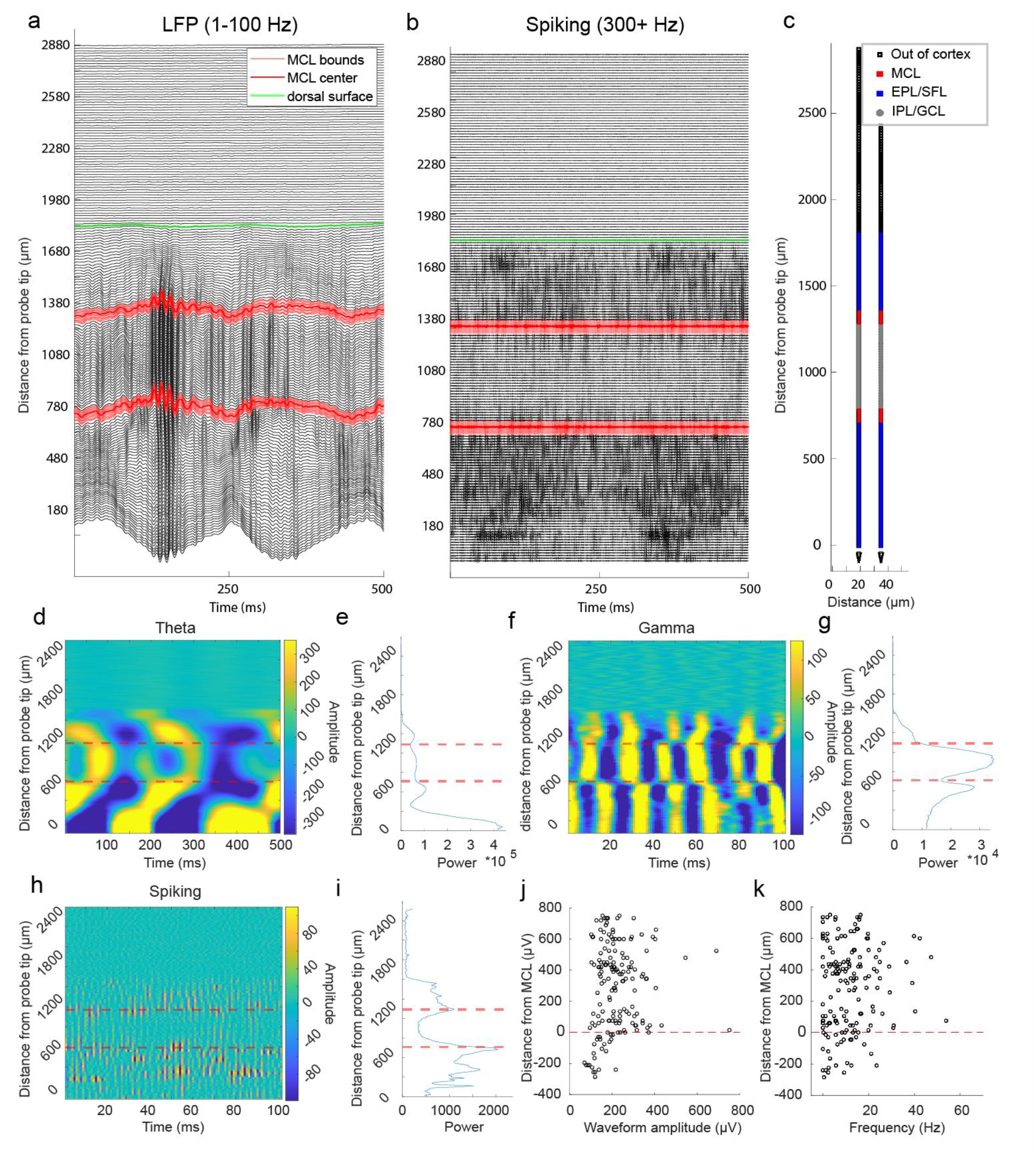
Single shank Neuropixels recording shows changes in LFP and spiking activity across the dorsal to ventral span of the OB. **(A)** Bandpass filtered LFP activity (1-100 Hz) from a single shank Neuropixels electrode array, (smoothed with 3.3m gaussian filter), covering ∼1.8 mm from the dorsal surface (green line) to the ventral OB. The estimated dorsal and ventral locations of the MCLs (mitral cell layers) are plotted (center of MCL dark red, bounds of MCL light red). The plot displays changes in LFP activity when moving from the dorsal surface (green line) towards the ventral OB. **(B)** A second plot of the same recording segment plotted in (a) is filtered for spiking activity (300-3000 Hz), showing dense clusters of spiking stretching from the dorsal and ventral superficial layers to the estimated MCLs. Notably, the central OB (deep layers, internal plexiform layer and granule cell layer) displays the least spiking across the bulb. **(C)** After MCL estimation, the resulting location of each recording site is labeled. **(D)** Theta LFP (2-10 Hz) amplitude extracted using Continuous Wavelet Transform based method and shows polarity reversal in theta frequency amplitude in the vicinity of the estimated MCLs. **(E)** Theta power is plotted one of the two columns of recording sites from the shank pictured in (c) showing local minima near the estimated MCLs. **(F-G)** Same as (d-e) is plotted for gamma amplitude (30-100 Hz) and shows a drop in amplitude near the MCL and a strong polarity reversal near the ventral MCL. **(H-I)** Same is plotted for spiking (300+ Hz) showing amplitude decreases rapidly when moving away from the MCL towards the IPL/GCL. **(J)** The waveform amplitude for each cell is plotted against its distance from the center of the MCL (positive distance indicates moving towards superficial cortex, negative distance indicates deeper OB). **(K)** The same is shown for the relationship between average spike rate and the distance from the MCL.

**Extended Data Figure 2:**
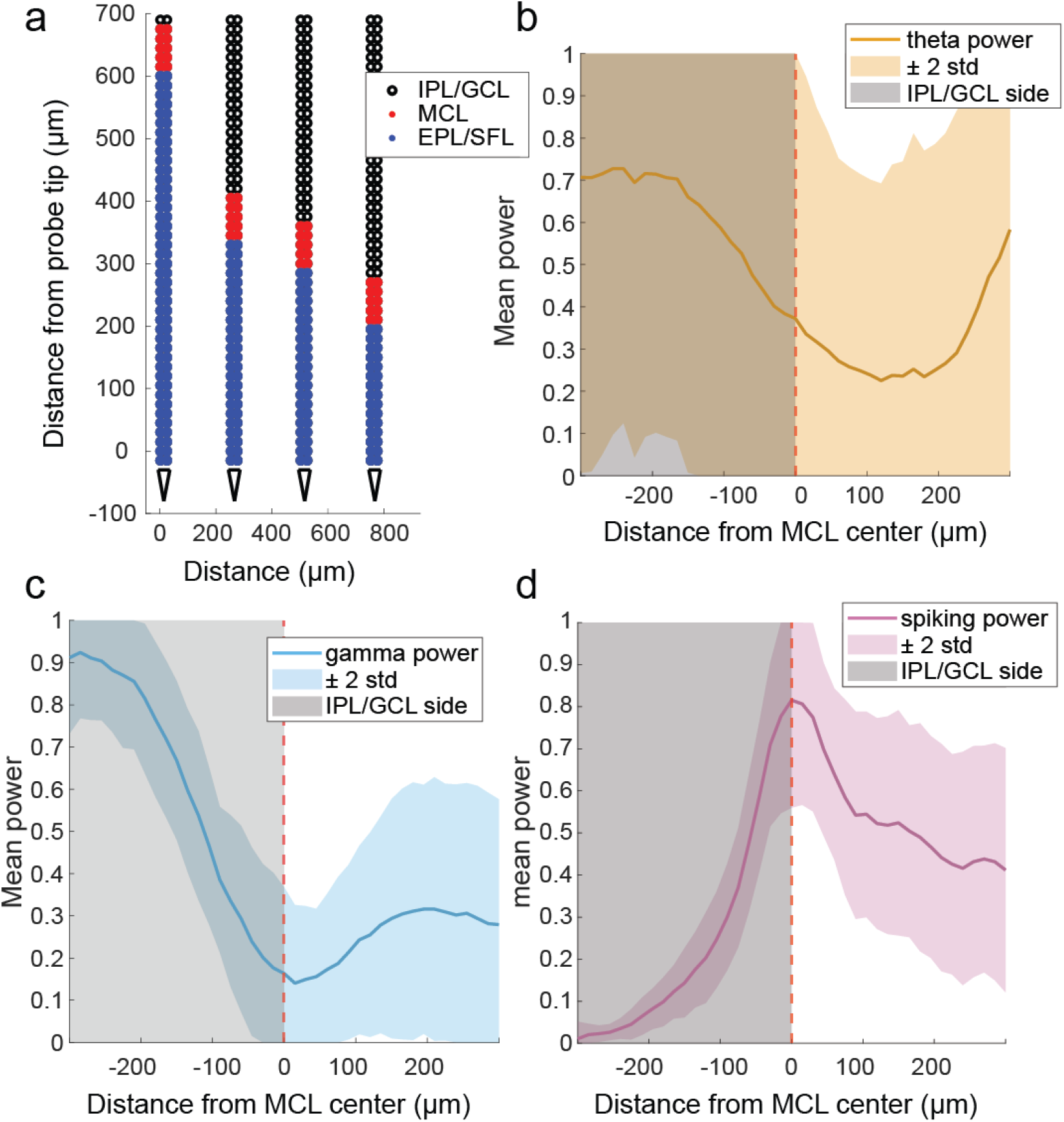
LFP changes relative to the estimated MCL. **(A)** A schematic for estimated layers across all four shanks from a single recording session of the ventral OB with the IPL/GCL (black), the MCL (red), and the EPL/SFL (blue) indicated. **(B)** Mean theta power (line) ±2 std (shading) across all shanks plotted as a function of distance from the MCL center shows that the exact point at which theta power drops off or reverses polarity relative to the estimated MCL (gamma drop off and spiking rise) is not reliable resulting in high variance for the drop in Theta power relative to the estimated center of the MCL. **(C)** The same is plotted but for gamma power. The drop off or polarity reversal of gamma power was more prominent and consistent across recordings and used to estimate MCL location. **(D)** The same is plotted for spiking power, showing a reliable rise in power when moving the GCL to EPL/SFLs, consistent with higher waveform amplitude and spiking rates of pyramidal cell activity in the OB as compared to axon-less granule cell in deep OB.

**Extended Data Figure 3:**
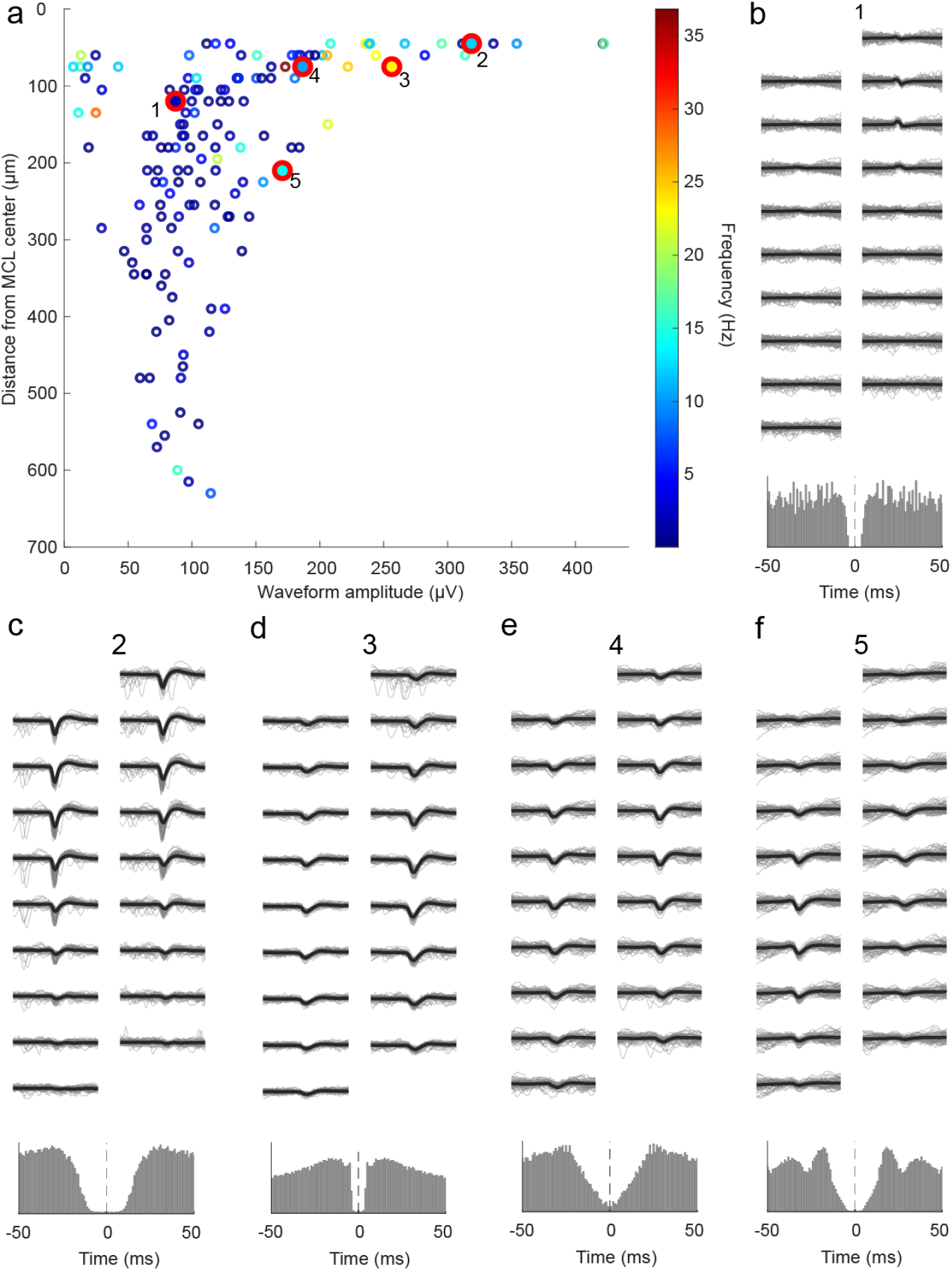
Clusters with waveforms in the IPL/GCL display low amplitude and low firing rates. **(A)** Distance from the MCL center (moving towards deeper OB) is plotted against waveform amplitude for all IPL/GCL (internal plexiform layer/ granule cell layer) clusters, showing a decline of both waveform amplitude and firing rate for clusters when moving away from the estimated MCL towards deeper OB. **(B)** (top) The mean waveform (black line) of an example cluster from the IPL/GCL area is plotted for 9 rows of recording sites (18 sites total). The mean waveform is plotted over 50 randomly selected individual waveforms (gray). Numbers for each cell correspond to the red highlighted clusters plotted in (a). (bottom) The auto-correlogram of the cluster (±50 ms) displays a refractory period. **(C-F)** The same is plotted for 4 additional IPL/GCL clusters. IPL/GCL clusters are shown to illustrate the physiological features of these clusters, but are not included in the analysis of cell or population activity.

**Extended Data Figure 4:**
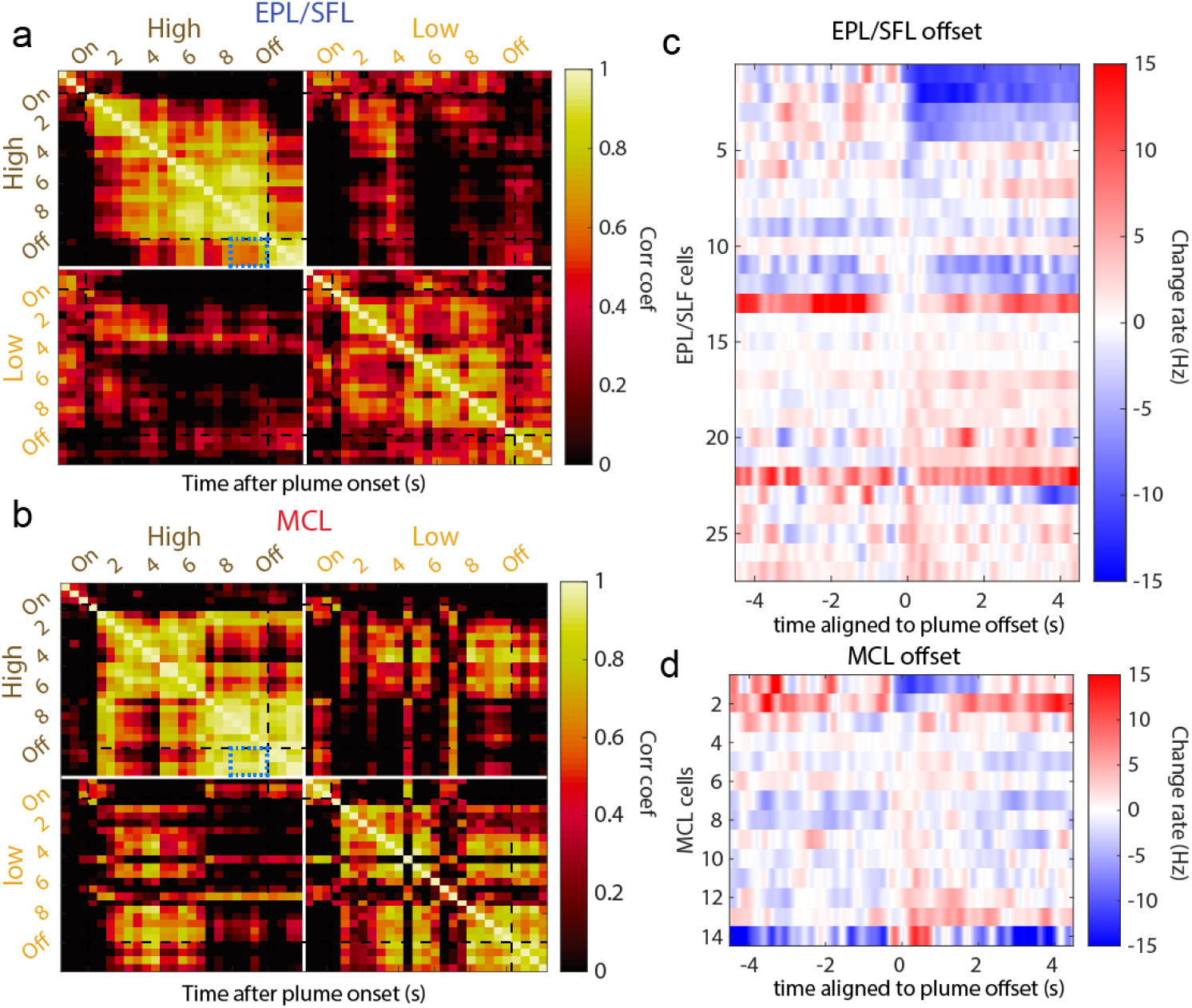
MC responses at the end of the plume correlated with responses after plume offset. **(A)** The average firing rate (Hz) of EPL/SFL cells exhibiting offset responses during high concentration trials (n=27 cells) is calculated for each 500ms time windows across the plume. Correlations between time windows are calculated within each concentration condition. The color axis is restricted to positive correlations for visualization purposes. **(B)** Same as (a) but for MCs with significant offset responses during high concentration trials (n=14 cells). Correlations between the last 2 seconds of the plume, and the first two seconds after plume offset (correlation coefficients that lie within the blue dotted square) are highly correlated for MCs (µ=.90, std=0.07), and significantly more correlated than they are for EPL/SFL cells (µ=.67, std=0.1) as determined by a Wilcoxon rank sum test (p<0.001, z=4.3908). This suggests MCs are highly likely to exhibit sustained offset responses. **(C)** For only EPL/SFL cells with significant offset responses compared to baseline activity before the plume (see methods, each trial is aligned to the end of the last whiff of each plume) offset response profiles are plotted. To visualize sustained versus rebounding offset responses within each cell type, each KDF is mean centered to show the change in firing rate between the end of the plume (-500 to 0 ms) and plume offset (0 to 500 ms). Therefore, all cells have significant offset responses relative to baseline, but as plotted here a sustained response will have no change in firing rate, and a rebounding response at offset would have a large change in firing rate. Reponses are sorted by the change in firing rate between these two time windows. **(D)** Same but for MCL cells with significant offset responses visualizing that in the 2 seconds following the plume, MCs are more likely to maintain activity after offset, as is shown by offset responses that are more similar to plume-elicited activity as compared to EPL/SFL cells.

**Extended Data Figure 5:**
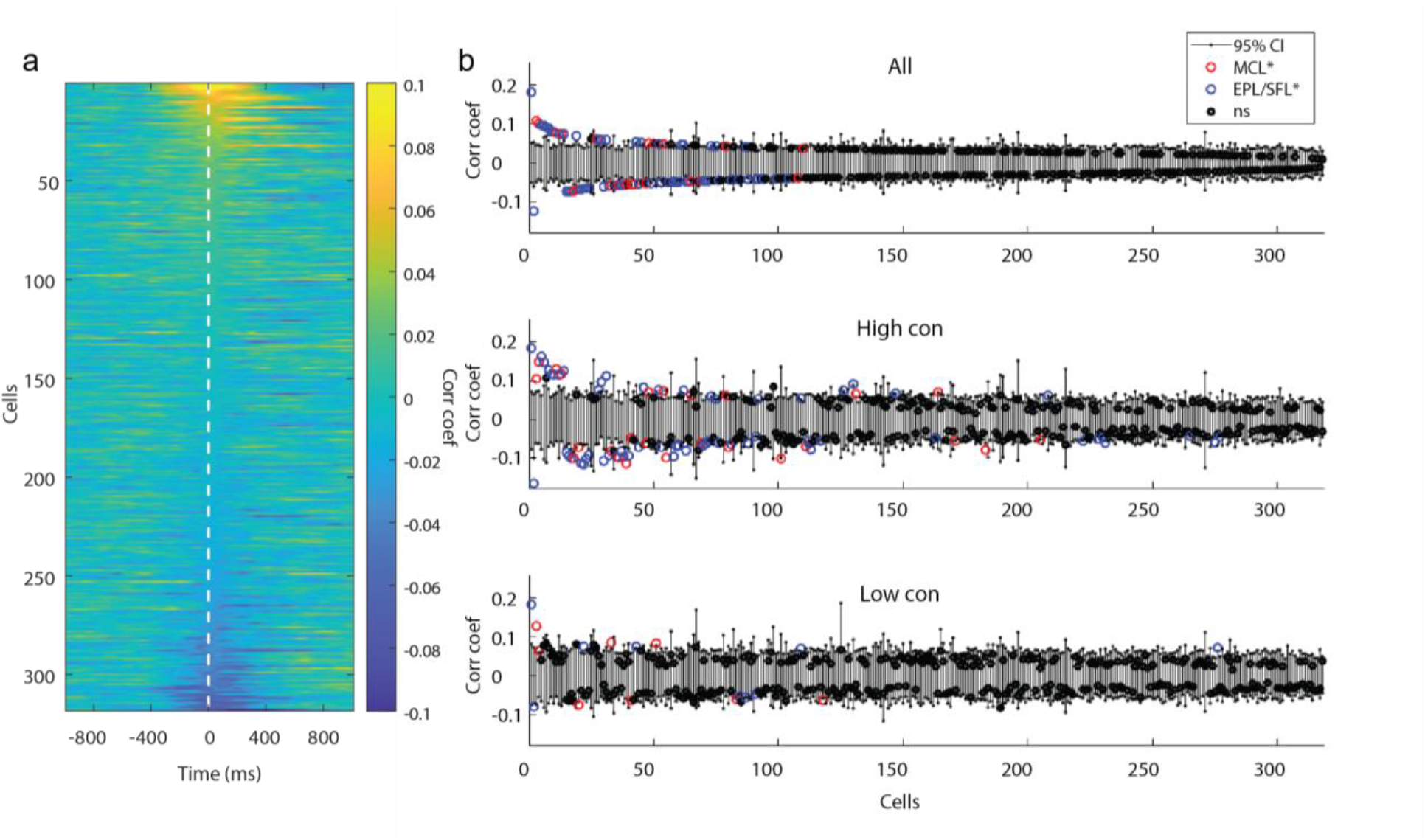
Binary correlation analysis indicated cells do not follow odor in a concentration invariant across the plume at the cellular level. **(A)** The cross-correlation between the binary odor signal and each cell’s spiking rate is calculated within each trial and then averaged across trials. Each row depicts the mean correlation coefficient between the cell’s spiking rate and the odor for each indicated lag ±1 s. Cells are sorted in order of decreasing magnitude of the max mean correlation coefficient within 0-500 ms lag. Color axis is restricted to -0.1-0.1 coefficient range for visualization purposes. **(B)** The mean correlation coefficients of all cells are plotted against their respective bootstrapped 95% confidence interval for all trials (top), for high concentration trials only (middle), and for low concentration trials only (bottom) and 23.9%, 26.7% and 5.4% (respectively) of cells were weakly but significantly correlated. Correlation coefficients are similar to those computed OB cell spiking and a non-binarized odor signal (Fig 6).

**Extended Data Figure 6:**
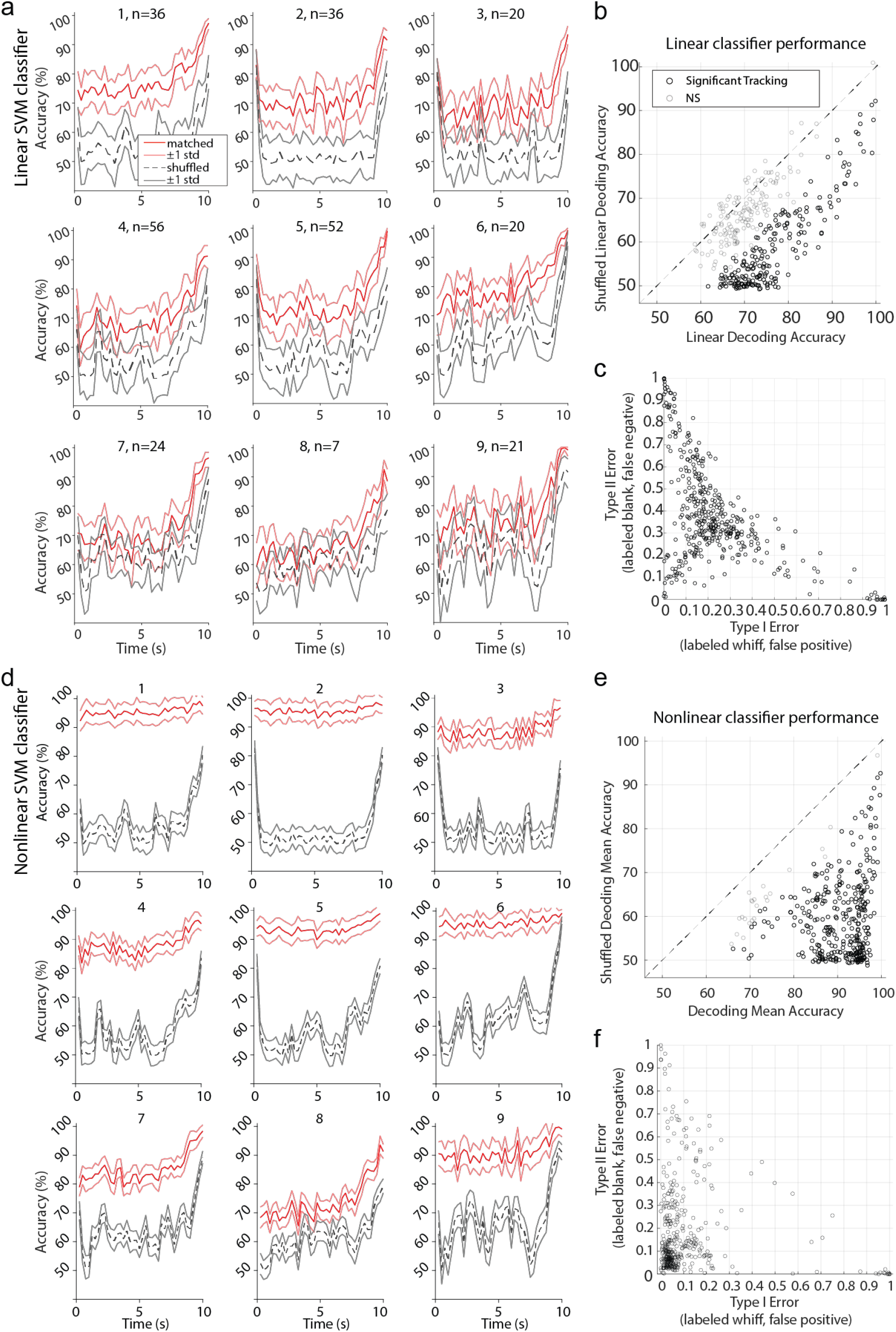
Radial Kernel SVM classifier is needed for robust decoding of whiffs and blanks. **(A)** Linear kernel SVM classifiers (red, mn±1std) run within each session show the majority of cell populations decode odor at above-chance level accuracy (grey, mn±1std) across the duration of the plume (250 ms binned windows from onset to offset). The number of cells in each session’s population is indicated above each plot. The binarized odor signal was biased towards odor being present at the start of the plume and biased towards odor being absent at the end of the plume as a byproduct of the alignment process (see methods: Decoding odor presence with nonlinear binary SCM classifiers), and as a result the shuffled chance level decoding performance increased at the beginning and end of the plume. **(B)** The mean accuracy for all time bins across all sessions are plotted against mean shuffled accuracy. If the mean accuracy did not exceed the shuffled confidence interval it is plotted in grey. **(C)** The mean Type I error, false positives where a timepoint during an odorless period is incorrectly classified as a whiff, and mean Type II error, false negatives where a timepoint during a whiff is incorrectly classified as a blank, are plotted for all classifiers from all time bins across sessions. **(D-F)** The same is plotted for radial kernel classifiers. Mean accuracy and mean error behavior (Fig 6e-f) are plotted again for comparison to (b-c), showing more robust classification across sessions and higher classification accuracy. Additionally, both linear and nonlinear classifiers tend to have a bias towards Type II Error.

